# Modular transcriptional responses to environmental changes

**DOI:** 10.1101/2023.01.09.523218

**Authors:** Marc Beringer, Bella Mattam, Rimjhim Roy Choudhury, Christian Parisod

## Abstract

Knowledge about the molecular underpinnings of phenotypic plasticity is still scarce and quantifying gene expression in response to abiotic and biotic factors enables to investigate transcriptional plasticity. RNAseq data on clones of the alpine plant *Biscutella laevigata* (Brassicaceae) subjected to control, cold, heat, drought and herbivory treatments assessed differentially expressed genes (DEGs) and transposable elements (DE-TEs) in comparison to similar experiments in *Arabidopsis thaliana*. Synergistic and trade-off DEGs presenting parallel and antagonistic regulation among treatments were further identified and used with networks of co-expressed DEGs to characterize transcriptional plasticity in response to environmental changes. Compared to *A. thaliana*, *B. laevigata* presented fewer DEGs that were mostly up-regulated by stronger expression shifts in response to environmental treatments. *Biscutella laevigata* showed constitutive expression of half of the *A. thaliana* DEGs. It further presented a higher proportion of synergistic DEGs, a lower number of trade-off DEGs and a transcriptome organized in environment-specific subnetworks. Several DE-TEs were identified as activated by heat and herbivory. The stress-tolerant perennial *B. laevigata* presents a highly modular transcriptional plasticity in response to environmental changes, contrasting with the more integrated transcriptome of *A. thaliana*.

**Significance statement:** Little is known about the molecular underpinnings of phenotypic plasticity. Here, focusing on expression shifts during changes in abiotic and biotic conditions, we highlight environment-responsive genes acting synergistically or antagonistically among treatments and underlying modular transcriptional plasticity in two Brassicacea species.

## Introduction

To survive and reproduce in changing environments, plants sense changes in abiotic and biotic factors, such as temperature, water availability and herbivores that trigger cascades of transcriptional changes leading to appropriate physiological responses (Kollist *et al*., 2019; Zhang *et al*., 2022). When the environmental stimulus impairs optimal growth and development, the stimulus becomes a stressor (Fujita *et al*., 2009). Despite numerous studies of gene expression under stress, our current understanding of transcriptional changes in response to environmental conditions represents mostly a post-hoc integration of genes characterized as triggered by a single factor (VanWallendael *et al*., 2019). The architecture of transcriptional responses to multiple environmental changes and thus the mechanisms of transcriptional plasticity remain elusive.

The transcriptome is anticipated to have evolved as a robust system, able to maintain functional interactions among gene products and integrity of the entire system against perturbations triggered by unpredictable environmental changes (Csete & Doyle, 2004; Kitano, 2004). Transcriptional plasticity in response to specific environmental triggers is likely key to maintain plant growth and reproduction, although it remains largely unknown to what extent environment-responsive genes act in synergy or are constrained by antagonistic interactions consistent with trade-offs (Stearns & Magwene, 2003; Lundgren & des Marais, 2020). It is thus necessary to assess patterns underlying transcriptional plasticity in response to independent environmental changes and identify possible functional modules of environment-responsive genes being regulated synergistically, or antagonistically (i.e. expression trade-offs).

Sensing of sub-optimal environments first induces a general stress response, leading to a more specific response (Zhang *et al*., 2022). The general response involves abscisic acid (ABA) and the expression of genes with ABA-responsive *cis*-elements (ABRE) as well as *dehydration-response element binding* (*DREB*)-type proteins (Nakashima *et al*., 2009) that act in synergy to regulate osmolyte levels and stomatal aperture, and to detoxify accumulating reactive oxygen species (Claeys & Inzé, 2013). Genes downstream of corresponding regulation cascades are expectedly constrained by expression trade-offs to trigger environment-specific responses (Kollist *et al*., 2019). Nevertheless, the transcriptional basis of physiological trade-offs such as the opening of stomata that mitigates heat stress and increases the risk of drought stress (Jacob *et al*., 2017), or defense against herbivores mediated by jasmonic acid (Howe & Jander, 2008) that also impedes growth (Devoto & Turner, 2005), deserves attention.

High-throughput sequencing of transcriptomes (RNAseq) is a powerful approach to assess expression and, in the absence of ample genomic resources (e.g. in non-model species), can be performed based on *de novo* transcriptome assemblies out of RNAseq reads themselves (Wang & Gribskov, 2017). Accordingly, the emergence of standardized practices for generating and analysing RNAseq data (Conesa *et al*., 2016) offers support towards meaningful conclusions on transcriptional plasticity under environmental changes. Here, we address transcriptional responses of *Biscutella laevigata*, a widespread species belonging to an early diverging Brassicaceae genus (Couvreur *et al*., 2010; Hendriks *et al*., 2022) to different environments. Being a textbook example of autopolyploidy linked to ice ages (Manton, 1937; Parisod & Besnard, 2007), diploids of *B. laevigata* occur across major ecological gradients (e.g. elevation from sea level to >2’000 m; Tremetsberger *et al*., 2002), or in extreme environments such as serpentine (Bürki *et al*., 2023) and mine soils (Babst-Kostecka *et al*., 2016). To investigate transcriptional plasticity, RNAseq on clones of a diploid *B. laevigata* subjected to cold, heat, drought and herbivory treatments, simulating abiotic and biotic stressors common to alpine environments, were compared with similar data from *Arabidposis thaliana* (Klepikova *et al*., 2016; Dubois *et al*., 2017; Nallu *et al*., 2018). This study thus aims at (i) evaluating approaches based on *de novo* transcriptome and genome assembly references to quantify gene expression, (ii) characterizing the transcriptional response of a non-model species in light of functional insights from a model plant to identify environment-responsive genes, and (iii) assessing patterns of plant transcriptional plasticity in response to environmental changes.

## Materials and Methods

### Plant material and environmental treatments

One diploid individual of *Biscutella laevigata* subsp. *austriaca* (Brassicaceae) was grown for a year, from seeds collected in the alpine population of Schneealpe (Steiermark, Austria; 1740 m above sea level; GPS: 47.6968°N, 15.6100°E), under standardized greenhouse conditions (16 h / 8 h light / dark, 22-26°C / 16-18°C, 65% relative air humidity, 20-40 kLux). Cuttings of that individual, including root and several leaves, were regenerated for ten weeks, forming ramets (i.e. clones) with at least eight new leaves.

To investigate transcriptional responses of *B. laevigata* to environmental treatments, individual clones were subjected to cold, heat, drought and herbivory conditions mimicking existing studies in *A. thaliana* (Klepikova *et al*., 2016; Dubois *et al*., 2017; Nallu *et al*., 2018). After an acclimation phase of seven days in a growth chamber under control conditions, the treatment phase started by stopping the watering of drought-treated clones until they showed first wilting leaves. At that moment the herbivory treatment (lasting 30 h) was started around noon, whereas the 24 h cold treatment was initiated at 6 pm and the heat treatment (9 h in total) was started at 9 am the next day. Accordingly, all treatments were terminated on the same day at 6 pm and treated leaves were harvested and snap frozen in liquid nitrogen. Subsequently relocated back to control conditions, all clones survived.

#### Control treatment

all clones were relocated to a growth chamber with cycles of 16 h of light (100-120 μM photosynthetically active radiation) and 8 h of dark at 22°C, under 45% relative air humidity and daily watering. Four clones were left under control conditions. Control conditions were similar for *A. thaliana*.

#### Cold treatment

four clones were relocated to a 4°C-room for 24 h under the same light cycle as in the control conditions, although with slightly lower light intensity (70-90 μM). *Arabidopsis thaliana* was treated the same (Klepikova *et al*., 2016).

#### Drought treatment

watering of three clones in the same growth chamber as the control treatment was stopped for a total of 11.5 days. Leaves were harvested 1.5 days after first evidence of wilting. In *A. thaliana*, watering was omitted for 3 days (Dubois *et al*., 2017).

#### Heat treatment

three clones were relocated to a growth chamber with the same settings as the control conditions. Temperature was raised gradually to 42°C during 3 h, where it remained for additional 6 h. A gradual increase in temperature has been shown to promote higher-fold transcriptome changes compared to sudden heat shocks (Larkindale & Vierling, 2008). Contrastingly, *A. thaliana* was subjected directly to 42°C for 6 h (Klepikova *et al*., 2016).

#### Herbivory treatment

three clones in the same growth chamber as control conditions were each applied eight larvae of the generalist herbivore Diamondback Moth *(Plutella xylostella;* 3^rd^ to 5^th^ instar; 6-9 mm in length, Ehlting *et al*., 2008) for 30 h. The mature leaf under treatment was encapsulated with a small plastic cage, wherein larvae were held (SI figure 2), resulting in between 10% and 20% of feeding damage by the time of harvest. In *A. thaliana*, 48 h old larvae of the Brassicacea specialist herbivore *Pieris rapae* were applied for 24 h (Nallu *et al*., 2018).

### RNA extraction and sequencing

Total RNA was extracted from 50-100mg of snap frozen leaf tissue of each experimental clone, using the RNAeasy mini kit (Qiagen) following a DNAse I treatment (Thermo Fisher Scientific). Samples were assessed using the Bioanalyzer 2100 system (Agilent) and considered for library preparation when the ribosomal RNA integrity number was over 7.0 (SI table 1).

In *B. laevigata* 17 RNAseq libraries (i.e. four biological replicates for the control and for the cold treatments, and three for the drought, heat and herbivory treatments) were processed at the next-generation sequencing platform of the University of Bern (Switzerland). Library preparation included the “TruSeq Stranded Total RNA with Ribo-Zero Plant”-kit (Illumina), ribosomal RNA depletion and size selection of 300 bp fragments. Sequencing on two lanes of an S2 flow cell of the NovaSeq6000 system yielded 50 bp paired-end (PE) reads (SI table 1). In *A. thaliana* 19 RNAseq libraries, sequenced as 50 bp single-end reads, were downloaded (SI table 2): two libraries each for cold, heat and their control treatments (Klepikova *et al*., 2016), two libraries for the drought treatment and three libraries for its control treatment (Dubois *et al*., 2017) and three libraries each for the herbivory and its control treatment (Nallu *et al*., 2018).

Adapter-trimmed raw PE-reads were quality controlled using FASTQC (Andrews, 2010) and summarized with MULTIQC (Ewels *et al*., 2016). To correct for sequencing errors, erroneous k-mers were identified and removed using the script *FilterUncorrectablePEfastq* in rCorrector (Song & Florea, 2015). An *in silico* rRNA-depletion was performed by mapping remaining reads to ribosomal RNA sequences from all organisms using bowtie2 (Langmead & Salzberg, 2012).

### De novo transcriptome assembly, annotation and quality control

To support a *de novo* transcriptome assembly, we further generated RNAseq libraries from seven tissues (leaf, senescent leaf, root, stem, closed flower bud, open flower and meristem) of a *B. laevigata* clone under control conditions (SI Figure 1b). Library preparation included the “TruSeq Stranded Total RNA Library Prep Human/Mouse/Rat”-kit (Illumina), ribosomal RNA depletion and a size selection step for 300 bp fragments. Sequencing on one lane of the HiSeq3000 system, yielded 150 bp PE reads (SI table 1).

Processed reads from these seven tissues and the reads of the largest libraries in each of the five treatments (SI table 1) were used as input for the *B. laevigata de novo* transcriptome. Assembly was conducted with TRINITY (Grabherr *et al*., 2011; Haas *et al*., 2013), with default parameters and the strandedness parameter set to -SS_lib_type FR. The *de novo* transcriptome was annotated with the Trinotate pipeline (Bryant *et al*., 2017) that identifies and aligns protein-coding sequences (CDS) to the swissprot protein database (561’568 protein sequences;www.uniprot.org; accessed in January 2020) using BLASTX with an e-value cut-off of 1e^-5^. The *de novo* assembled transcripts were further linked with gene annotations from the genome assembly of *B. laevigata* and *A. thaliana* through a BLASTN with an e-value cut-off of 1e^-10^.

The *de novo* transcriptome assembly was quality-controlled by assessing the proportion of input reads used in the final assembly, as well as the mapping of the twelve input RNAseq libraries to the transcriptome using bowtie2, allowing for up to 20 valid alignments per read. We measured the N50 value and gauged completeness of the *de novo* transcriptome of *B. laevigata* by conducing BLASTX with the *A. thaliana* protein database (27’465 protein sequences; UniProt accession number UP000006548), using an e-value cut-off of 1e^-20^ and only returning the first hit. We computed the BUSCO-score (Benchmarking Universal Single-Copy Orthologs, v4.1.4; Manni *et al*., 2021), using the “eudicotyledons_odb10” database (Creation date: 2019-11-20, with 2326 BUSCOs).

### Gene expression quantification and differential gene expression analysis

Gene expression among the 17 libraries encompassing environmental treatments in *B. laevigata* was quantified based on the *de novo* transcriptome CDS, as well as the CDS of the *B. laevigata* genome assembly (comprising of 54’457 genes and 88’133 isoforms; table 1), enabling comparisons of transcriptome-based and genome-based quantification as well as direct comparisons with genome-based analyses in *A. thaliana* (Araport11_genes.201606.cds.fasta.gz; Cheng *et al*., 2017).

**Table 1.**
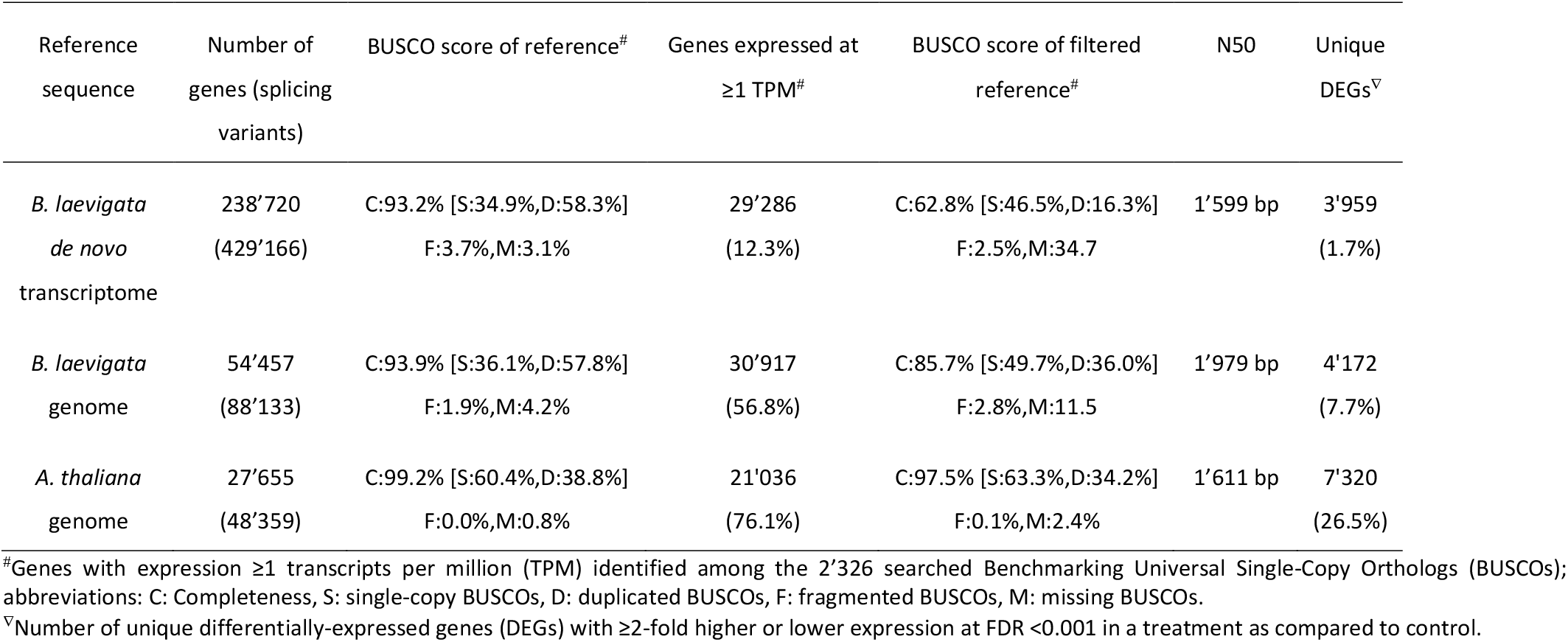
Quality metrics of the three sequence references used to quantify gene expression under environmental treatments

| Reference<br>sequence | Number of<br>genes (splicing<br>variants) | BUSCO score of reference<sup>#</sup> | Genes expressed at<br>≥1 TPM<sup>#</sup> | BUSCO score of filtered<br>reference<sup>#</sup> | N50 | Unique<br>DEGs<sup>▽</sup> |
| --- | --- | --- | --- | --- | --- | --- |
| *B. laevigata*<br>*de novo*<br>transcriptome | 238'720<br>(429'166) | C:93.2% [S:34.9%,D:58.3%]<br>F:3.7%,M:3.1% | 29'286<br>(12.3%) | C:62.8% [S:46.5%,D:16.3%]<br>F:2.5%,M:34.7 | 1'599 bp | 3'959<br>(1.7%) |
| *B. laevigata*<br>genome | 54'457<br>(88'133) | C:93.9% [S:36.1%,D:57.8%]<br>F:1.9%,M:4.2% | 30'917<br>(56.8%) | C:85.7% [S:49.7%,D:36.0%]<br>F:2.8%,M:11.5 | 1'979 bp | 4'172<br>(7.7%) |
| *A. thaliana*<br>genome | 27'655<br>(48'359) | C:99.2% [S:60.4%,D:38.8%]<br>F:0.0%,M:0.8% | 21'036<br>(76.1%) | C:97.5% [S:63.3%,D:34.2%]<br>F:0.1%,M:2.4% | 1'611 bp | 7'320<br>(26.5%) |

<sup>#</sup>Genes with expression ≥1 transcripts per million (TPM) identified among the 2'326 searched Benchmarking Universal Single-Copy Orthologs (BUSCOs); abbreviations: C: Completeness, S: single-copy BUSCOs, D: duplicated BUSCOs, F: fragmented BUSCOs, M: missing BUSCOs.
<sup>▽</sup>Number of unique differentially-expressed genes (DEGs) with ≥2-fold higher or lower expression at FDR <0.001 in a treatment as compared to control.

Gene expression was quantified using RSEM estimated counts (Li & Dewey, 2011) that served as input for differential gene expression analyses using edgeR (Robinson *et al*., 2009). Following a within-sample normalization, gene expression was evaluated as transcript per million (TPM) to ensure independence from transcript length and increase comparability among samples (Li *et al*., 2009; Wagner *et al*., 2012). To further account for RNAseq library sizes, cross-sample normalization of TPM expression was performed with the “trimmed mean of M values” (TMM) method (Robinson & Oshlack, 2010). Differential gene expression analyses were conducted on RSEM estimated counts using the TRINITY script *run_DE_analysis.pl* to estimate log_2_-fold-change (logFC) and *analyse_diff_expr.pl* to extract ≥2-fold differentially expressed genes (DEGs) at a Benjamini-Hochberg false discovery rate (FDR) <0.001 (Benjamini & Hochberg, 1995).

### Gene functional enrichment analysis

To what extent sets of genes (e.g. treatment-specific DEGs) were enriched in particular functional categories was assessed through gene ontology (GO) enrichment analyses, using the topGO R-package (Alexa *et al*., 2006). We used GO-term annotation files from the Trinotate-pipeline for the *de novo* transcriptome and genome assembly of *B. laevigata* and the latest Araport11 annotation of Berardini *et al*. (2004) for *A. thaliana*(https://www.arabidopsis.org; downloaded in November 2021). Although overrepresented GO-terms should be interpreted with caution, functional insights gathered here for *B. laevigata* appear particularly credible, given relatedness with the model plant *A. thaliana* that benefits from ample experimental support (Primmer *et al*., 2013).

### Weighted gene co-expression network analysis

Genome-based expression data were analysed through weighted gene co-expression networks, using the WGCNA R-package (Langfelder & Horvath, 2008), to assess the transcriptome architecture. Nodes (genes) were connected through edges, weighted between 0 and 1 based on the correlation of gene expression profiles (i.e. “adjacency”) using TMM-normalized gene expression values from environmental treatments. Genes showing similar expression profiles across samples were grouped into modules that were correlated to environmental treatments.

Independent co-expression networks were generated for *B. laevigata* and *A. thaliana* following the pipeline available at https://github.com/Persilian/WGCNA/wiki. Briefly, using TMM-normalized expression matrices with an optimal soft-threshold power set to 11 and fitting the scale-free topology criterion as r^2^ = 0.95 for the *Biscutella* data and r^2^ = 0.89 for the *A. thaliana* data, networks were computed with the function “blockwiseModules()” using corType = “pearson”, networkType = “unsigned”, maxBlockSize = 10000, TOMType = “unsigned”, minModuleSize = 30 and mergeCutHeight = 0.1. Only DEGs were visualized into subnetworks using *Cytoscape* (Shannon *et al*., 2003) by subsetting the adjacency matrices containing pairwise edge-weights among all genes. The “Perfuse Force Directed Layout” was applied (1000 iterations, spring coefficient = 0.5, spring length = 20, node mass = 1000 and “Force deterministic layouts”) and only edges above a threshold (adj_threshold = 0.5 for *B. laevigata* and 0.7 for *A. thaliana)* were visualized. A higher edge-weight threshold was selected for *A. thaliana* because a threshold ≥0.5 included over 200’000 edges and precluded subsequent analyses.

### Quantification of transposable element expression

The expression of 8’646 full-length sequences of transposable elements (TEs) annotated in the *B. laevigata* genome was quantified using the SalmonTE pipeline (Jeong *et al*., 2018). Briefly, a fasta file containing internal coding sequences of TEs extracted using BEDtools-getfasta (Quinlan & Hall, 2010) was used for rapid quasi-mapping of RNAseq reads from the 17 libraries of environmental treatments, resulting in TE expression in TPM. Differential expression among environmental treatments was tested using a Generalized Linear Model, for each TE copy as well as their aggregation into TE clades.

To distinguish differentially expressed TEs (DE-TE) whose expression is potentially controlled directly by an environmental treatment from DE-TEs showing co-expression with adjacent genes responding to the treatment, reverse-transcriptase sequences from all TEs were extracted using TEsorter, aligned using MAFFT (Katoh *et al*., 2002) and clustered using maximum-likelihood phylogenetic trees in iTOL (Letunic & Bork, 2021). Monophyletic clades of DE-TE copies being consistently up-regulated in response to the same environmental treatment were accordingly identified as environment-activated and characterized through homology search using CENSOR in Repbase (Kohany *et al*., 2006).

## Results

### Gene expression based on de novo transcriptome vs genome assembly

We sequenced a total of 1489.6M raw sequencing reads including transcripts of *B. laevigata* expressed in seven tissues under control conditions (ENA accession PRJEB48599; ca. 386M PE-reads) and in 17 replicated samples subjected to environmental treatments (ENA accession PRJEB48469; ca. 1103M PE-reads). After removal of 2.4% reads with putative sequencing errors and 35.1% of reads mapping to ribosomal RNA sequences across all RNAseq libraries, processed libraries contained between 11 and 120M PE-reads (average of 37.2M reads per library; SI table 1).

The *de novo* transcriptome presented 238’720 genes and 429’166 isoforms (table 1), corresponding to 16’397 protein sequences with ≥80% homology on swissprot, whereas 17’460 genes with ≥80% homology to *A. thaliana* proteins were identified. Similar N50 values of CDS were achieved for the *de novo* transcriptome (1’599 bp), the *B. laevigata* genome assembly (1’979 bp) and the Araport11 genome assembly of *A. thaliana* (1’611 bp). Completeness scores of BUSCO across references was over 93% (table 1) and most (97.8%) input reads mapped back to the *de novo* transcriptome. The 17 libraries of environmental treatments mapped on average at 72.8% to the *de novo* transcriptome (SI table 3), and at 38.6% to the *B. laevigata* genome assembly (SI table 4). The *A. thaliana* RNAseq data (19 libraries) mapped on average at 71.6% to the Araport11 CDS (SI table 5).

Of the 238’720 transcripts assembled in the *de novo* transcriptome of *B. laevigata*, 29’286 (12.3%) were expressed at ≥1 TPM. Similarly, the *B. laevigata* genome assembly presented 30’917 genes as expressed at ≥1 TPM (56.8% of the 54’457 genes), supporting convergent insights to be gathered from RNAseq, independent of the reference sequence. In *A. thaliana*, 21’036 genes were expressed at ≥1 TPM (76.1% of the 27’655 genes in the Araport11 reference; table 1).

### Differential gene expression in response to environmental treatments

Transcriptional responses to cold, heat, drought and herbivory were characterized and compared between *B. laevigata* and *A. thaliana* by the number of DEGs, the proportion of DEGs exclusive to a treatment, the number of synergistic DEGs (up-regulated in more than one treatment) and of antagonistic (i.e. trade-off) DEGs that are up- and down-regulated among treatments, as well as the strength of expression shifts (logFC).

Considering only genes presenting at least a two-fold shift in their expression (logFC ≥1, FDR <0.001) in response to a treatment as DEGs, 3’959 (13.5% of transcripts expressed ≥1 TPM) and 4’172 (13.5% of genes expressed ≥1 TPM) unique DEGs were identified in the transcriptome- and genome-based analyses of *B. laevigata*, respectively (figure 1 and SI table 6). Analyses based on the *de novo* transcriptome performed adequately, although identifying only 2’615 (62.7%) of the 4’172 DEGs identified using the genome-based approach. *Arabidopsis thaliana* revealed a larger proportion of DEGs than *B. laevigata*, with 7’320 unique DEGs (26.5% of all annotated genes; SI table 7).

**Figure 1.**
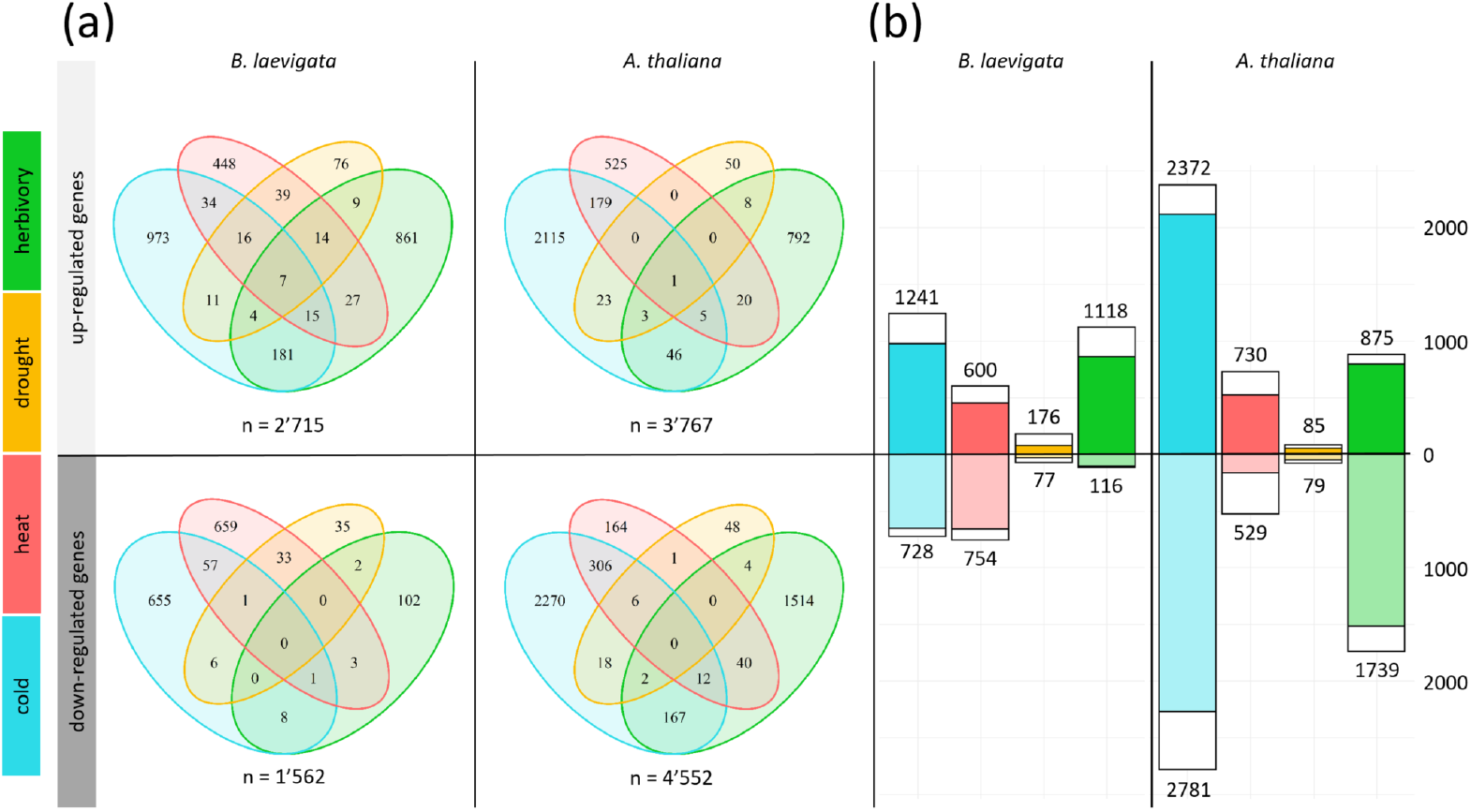
Differentially expressed genes (DEGs, with ≥2-fold change) in *Biscutella laevigata* and *Arabidopsis thaliana* in response to cold, heat, drought and herbivory treatments. **a)** Venn diagrams showing the number of up- and down-regulated genes in response to the different environmental treatments. **b)** Numbers of up- and down-regulated DEGs per treatment, with treatment-specific DEGs presented by the corresponding colour and DEGs in response to multiple treatments shown in white.

The *B. laevigata* and *A. thaliana* DEG sets had 1’504 DEGs in common, likely representing a core set of environment-responsive genes in Brassicaceae. Noticeably, 3’983 of the DEGs identified in *A. thaliana* (i.e. 55% of its DEGs) presented constitutive expression above ≥1 TPM in *B. laevigata*, while 1’000 more were non-expressed (<1TPM), suggesting that the alpine *B. laevigata* evolved constitutive expression of environment-responsive genes since divergence from the common ancestor with *A. thaliana*.

Contrasting patterns of expression changes between species were further apparent from the majority of genes being up-regulated (2’715; 5%) and only 1’562 genes (2.9%) being down-regulated in *B. laevigata*, whereas *A. thaliana* presented an opposite pattern with fewer up- (3’767; 13.6%) than down-regulated (4’552; 16.5%) genes (two-proportions Z-tests p-values <0.001; figure 1a). Most up-DEGs in *B. laevigata* responded to cold and herbivory, sharply contrasting with the majority of DEGs being down-regulated under these treatments in *A. thaliana* (figure 1b). In both species, cold and herbivory triggered the most treatment-specific sets of genes (>80% of cold- and herbivory-DEGs exclusive to those treatments), whereas the heat treatment shared a considerable number of DEGs with the response to cold. Drought triggered more treatment-unspecific genes in *B. laevigata* and *A. thaliana*, highlighting only 44% and 60% of exclusive DEGs respectively and indicating a pivotal role of genes involved in water homeostasis under our environmental treatments.

Transcriptional synergies across treatments were evaluated by proportions of identified synergistic DEGs. In *B. laevigata* seven genes were up-regulated in response to all environmental treatments, while *A. thaliana* presented only one such DEG (see Supplementary Text T1). Furthermore, *B. laevigata* presented more synergistic DEGs than *A. thaliana*, with 357 DEGs (8.5% of all DEGs) and 285 DEGs (3.9%; two-proportions Z-test p-value <0.001), respectively.

Contrastingly, *B. laevigata* presented 105 (2.5%) trade-off DEGs, representing a significantly lower proportion than in *A. thaliana* (999 out of 7’320, 13.6%; two-proportions Z-test, p-value <0.001). The 105 trade-off DEGs in *B. laevigata* were involved in a total of 215 expression trade-offs across treatments, whereas the 999 genes of *A. thaliana* were involved in 2’187 trade-offs. Most identified trade-off DEGs were involved in cold and herbivory responses, with 50 out of 215 cases (23.3%) in *B. laevigata* and 788 cases (36.0%) in *A. thaliana*, whereas cold and heat responses further highlighted 23 (10.7%) and 110 (5.0%) cases, respectively. The drought treatment triggered few trade-off DEGs in both species, with 9 and 70 for *B. laevigata* and *A. thaliana*, respectively.

Expression shifts of DEGs (SI figure 3) were significantly stronger in *B. laevigata* (average logFC = 3.21) compared to *A. thaliana* (average logFC = 2.55; Wilcoxon paired test p-value <0.001). Despite the relatively few drought DEGs in *B. laevigata*, these genes achieved the strongest shifts in expression, with an average logFC of 5.21 for drought up- and of −4.98 for down-regulated genes. Such >30-fold change in the expression of drought-responding genes in *B. laevigata* strongly contrasted the drought response of *A. thaliana* that shifted expression the weakest under this treatment (logFCs of 1.58 and −1.61 for up- and down-regulated genes, respectively).

### Differential expression of transposable elements in response to environmental treatments

Out of 8’646 full-length TE sequences in the *B. laevigata* genome, 299 TE copies presented differential expression (DE-TEs; logFC ≥1, FDR <0.001) in response to cold, heat, drought or herbivory. The majority of DE-TEs (210) were up-regulated in response to a specific treatment (figure 2a). Environmentally triggered DE-TEs were distinguished from DE-TE copies being expressed following changes in expression of nearby genes, through the identification of clades presenting only related DE-TE copies responding to the same treatment. Two clades of LTR-*Copia* presented an expression pattern consistent with transcriptional activation by a specific environmental trigger (figure 2b). One of these TE clades presented clear up-regulation of seven copies in response to the heat treatment and showed strong homology to *TERESTRA* from multiple Brassicaceae. Another DE-TE clade presented consistent up-regulation of 27 copies under herbivory and included nine related copies showing clear homology to *Atcopia31* and 18 copies showing homology to *Atcopia93* (also called EVD). As genes located closest up- or down-stream of these DE-TEs showed no indication of co-expression (SI table 8), which suggests limited interactions between genes and TEs.

**Figure 2.**
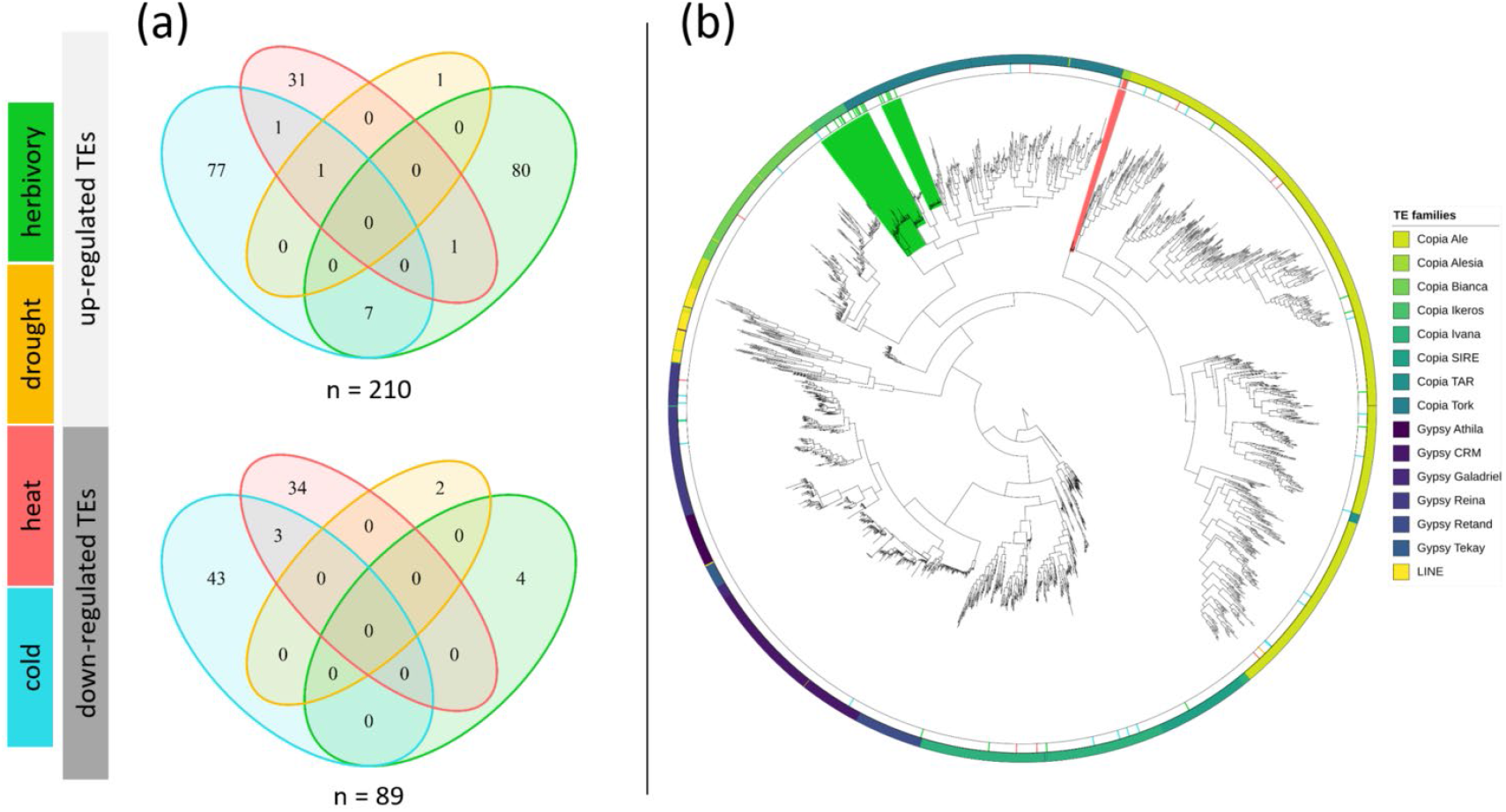
Differentially expressed transposable elements (DE-TEs) in *Biscutella laevigata* in response to cold, heat, herbivory and drought treatments. **a)** Venn diagrams showing the number of full-length TE copies showing differential expression in response to environmental treatments in *B. laevigata*. **b)** Phylogenetic tree of TE copies based on their reverse-transcriptase sequences and delineating main TE lineages of long terminal repeat retrotransposons Copia and Gypsy, and long interspersed nuclear element (LINE) according to the legend on the right. Up-regulated DE-TEs in response to environmental treatments are mapped on the inner annotation circle, with clades of consistently up-regulated DE-TEs coloured in green for herbivory and red for heat.

### Functional insights on genes controlled by environmental treatments

As expected under environmental stress, all transcriptional responses to treatments were enriched in DEGs related to ABA, water homeostasis and oxidative stress in both *B. laevigata* and *A. thaliana* (SI tables 9-24).

Cold up-regulated genes in *B. laevigata* were chiefly involved in the “regulation of metabolic process” (222 DEGs) and “defense response” (201 DEGs). In particular, the cold response involved up-regulation of 82 genes annotated as “defense response to bacterium”, 40 genes annotated as “defense response to fungus”. On the other hand, down-regulation of genes annotated with “response to insect” (6 DEGs) and “response to herbivore” (5 DEGs) indicated a possible trade-off between cold response and defense against herbivores, but not against pathogens. In contrast, *A. thaliana* appeared to down-regulate pathogen defense in response to cold, as indicated by 75 down-DEGs related to “defense response to bacterium” and 47 down-DEGs annotated as “defense response to fungus”. In both species, the cold treatment induced the up-regulation of *STCH4* (AT2G24500) that is known to confer cold tolerance through increased translation rates of *CBF* proteins and the accumulation of *CBF1-3* proteins (i.e. *DREB1b, DREB1c* and *DREB1a)*. Furthermore, in both species the circadian clock genes *LHY* (AT1G01060) and *CCA1* (AT2G46830) were up-regulated under cold conditions. Starch metabolism also appeared affected by the cold treatments, as indicated by the up-regulation of the β-amylase genes *BAM1* (AT3G23920) and *BAM3* (AT4G17090).

Heat induced the up-regulation of genes involved in “protein folding” and “response to salt stress” GO-terms in both species (respectively, 60 and 50 DEGs in *B. laevigata* and 40 and 34 DEGs in *A. thaliana)*.In *B. laevigata*, genes involved in “response to osmotic stress” (60 DEGs) and “response to ABA” (51 DEGs) were up-regulated, while 11 DEGs involved in “photosynthesis” were down-regulated. In contrast, *A. thaliana* down-regulated “response to ABA” (17 DEGs) and up-regulated “photosynthesis” (5 DEGs). In both species, the chloroplast-localized *HSP21* (AT4G27670), the mitochondrion-localized *HSP23* (AT4G25200), the cytosolic *HSP90.1* (AT5G52640), *HSFA2* (AT2G26150) and *HSFA3* (AT5G03720). Similar to the cold response, the extreme heat treatment induced up-regulation of *BAM1* in both species, indicating that mobilization of starch may be important during temperature stress.

In both *B. laevigata* and *A. thaliana*, the drought response was enriched in DEGs related to “response to water deprivation” (27 and 5 DEGs) and “response to ABA” (27 and 9 DEGs) and, noticeably, “response to cold” (21 and 7 DEGs). Furthermore, *B. laevigata* up-regulated 24 genes responding to osmotic stress and down-regulated genes involved in “cell wall organization” (11 DEGs) and “pectin catabolic process” (4 DEGs), suggesting an impact on growth. On the other hand, *A. thaliana* up-regulated five genes involved in light harvesting of photosystem I and II, indicating short-term increase in photosynthesis under drought. Four genes were commonly drought up-regulated in both species, including the *highly ABA-induced PP2C gene 2* (*AIP1*; AT1G07430) involved in stomatal aperture, *PUB19* (AT1G60190, a U-Box E3 ubiquitin ligase), *AFP1* (AT1G69260, an ABA-involved transcription factor) and *AITR1* (AT3G27250, ABA-induced transcriptional repressor).

In both species, the herbivory treatment mainly induced genes involved in “defense response”, although considerably more numerous in *B. laevigata* (210 up-DEGs) than in *A. thaliana* (19 up-DEGs). Similarly, *B. laevigata* up-regulated more genes than *A. thaliana* involved in “signal transduction” (145 and 37 DEGs, respectively), “response to jasmonic acid” (67 and 13) and “glucosinolate metabolic process” (37 and 3). Noticeably, down-regulation of genes involved in “cold-acclimation” and “response to cold” was apparent in both species, suggesting a general trade-off between herbivory and cold stresses. Indicative of a differential impact on growth, *B. laevigata* up-regulated 82 genes responding to ABA, whereas *A. thaliana* down-regulated 63 of them. More specifically, herbivory induced up-regulation of *jasmonate-induced oxygenase 3* (*JAO3*, AT3G55970) in both species, although *P. xylostella* feeding on *B. laevigata* further induced the up-regulation of homologues of *terpene synthase 3* (*TPS03*, AT4G16740), which was down-regulated in *A. thaliana*. Similarly, *CYP79B2* (AT4G39950), a key enzyme of the glucosinolate biosynthesis, was identified as up-regulated under herbivory in *B. laevigata*.

The 1’504 DEGs shared among *B. laevigata* and *A. thaliana* presented enrichment in transcription factors (SI table 25), with 66 such putative environment-responsive master regulators identified in both species (SI table 26). They were related to growth (e.g. the *growth regulating factor 3* and *4;*AT2G36400 and AT3G52910 respectively) and stress-related ethylene response factors (i.e. members of the *DREB* sub-family; AT1G12610, AT1G19210; AT1G64380; AT5G21960) as well as regulators of flowering and the circadian rhythm (e.g. *FLM and LHY;* AT1G77080 and AT1G01060, respectively).

In *B. laevigata*, the 357 synergistic DEGs were enriched in GO-terms related to water homeostasis and oxidative stress. Enrichment in DEGs related to “response to water deprivation” (45 DEGs), “response to abscisic acid” (43 DEGs), “response to salt stress” (36 DEGs) and “response to oxidative stress” (26 DEGs; SI table 27) supported a general role of drought related genes in all our treatments. To a lesser extent, this was also the case for the 285 synergistic DEGs in *A. thaliana*, with enrichment of processes such as response to salt stress” (18 DEGs) and “response to oxidative stress” (15 DEGs; SI table 28)

Trade-off DEGs in *B. laevigata* were chiefly involved in biological processes such as “oxidation-reduction process” (15 DEGs), “carbohydrate metabolic process” (12 DEGs) and “response to water deprivation” (7 DEGs; SI table 29). Although *A. thaliana* trade-off DEGs were enriched in “response to cold” (52 DEGs), “response to ABA” (44 DEGs), they shared processes such as “oxidation-reduction process” (41 DEGs) and “response to water deprivation” (31 DEGs; SI table 30).

Among the numerous species-specific DEGs, the 2’158 *B. laevigata-specific* DEGs were mostly enriched in defense-, stress-, ABA-, as well as glucosinolate biosynthesis-related GO-terms (SI table 31), whereas the 5’816 DEGs specific to *A. thaliana* appeared to be predominantly enriched in terms related to “regulation of transcription, DNA-templated” and “translation” (SI table 32).

A considerable number of DEGs specific to *A. thaliana* (4’983) did not present differential expression in *B. laevigata*, with a majority of them (3’983) presenting constitutive expression ≥1 TPM in that species. Enrichment in functions such as “response to ABA”, “response to cold”, “response to heat” and “response to water deprivation” suggested a persistent basal expression of abiotic stress-related genes in the alpine species. Consistent with the slow growth and perennial life strategy of *B. laevigata*,these constitutively-expressed genes were involved in energy intensive processes such as “protein phosphorylation”, “photosynthesis” and “circadian rhythm” (153, 35 and 33 genes respectively; SI table 33).

### Transcriptional architecture of responses to environmental changes

Considering genes showing substantial expression variance across samples of *B. laevigata* (41’613 genes out of 54’457) and *A. thaliana* (26’051 out of 27’655), weighted gene co-expression network analysis grouped them into modules based on their correlated expression profiles across environmental treatments. Although the *B. laevigata* co-expression network encompassed considerably more genes, expression profiles resolved into 78 modules containing between 35 and 2’440 genes (SI table 34), whereas *A. thaliana* presented 93 modules containing between 36 and 3’002 genes (SI table 35). Networks of the most strongly connected DEGs (i.e. DEG-networks) included 2’625 DEGs in *B. laevigata* and 4’465 DEGs in *A. thaliana* (figure 3). The DEG-network in *B. laevigata* delineated four distinct subnetworks of up-regulated DEGs specific to either cold, heat, drought or herbivory responses and a fifth subnetwork consisting of mostly down-regulated DEGs (SI figure 4). In contrast to such a modular organisation of DEGs in *B. laevigata*, the DEG-network of *A. thaliana* presented overall higher correlations across gene expression profiles and a diffuse distribution of treatment-specific up-DEGs (SI figure 5). Together with the considerable number of *A. thaliana* DEGs being constitutively expressed in *B. laevigata*, this suggest a differential architecture of transcriptional plasticity between species.

**Figure 3.**
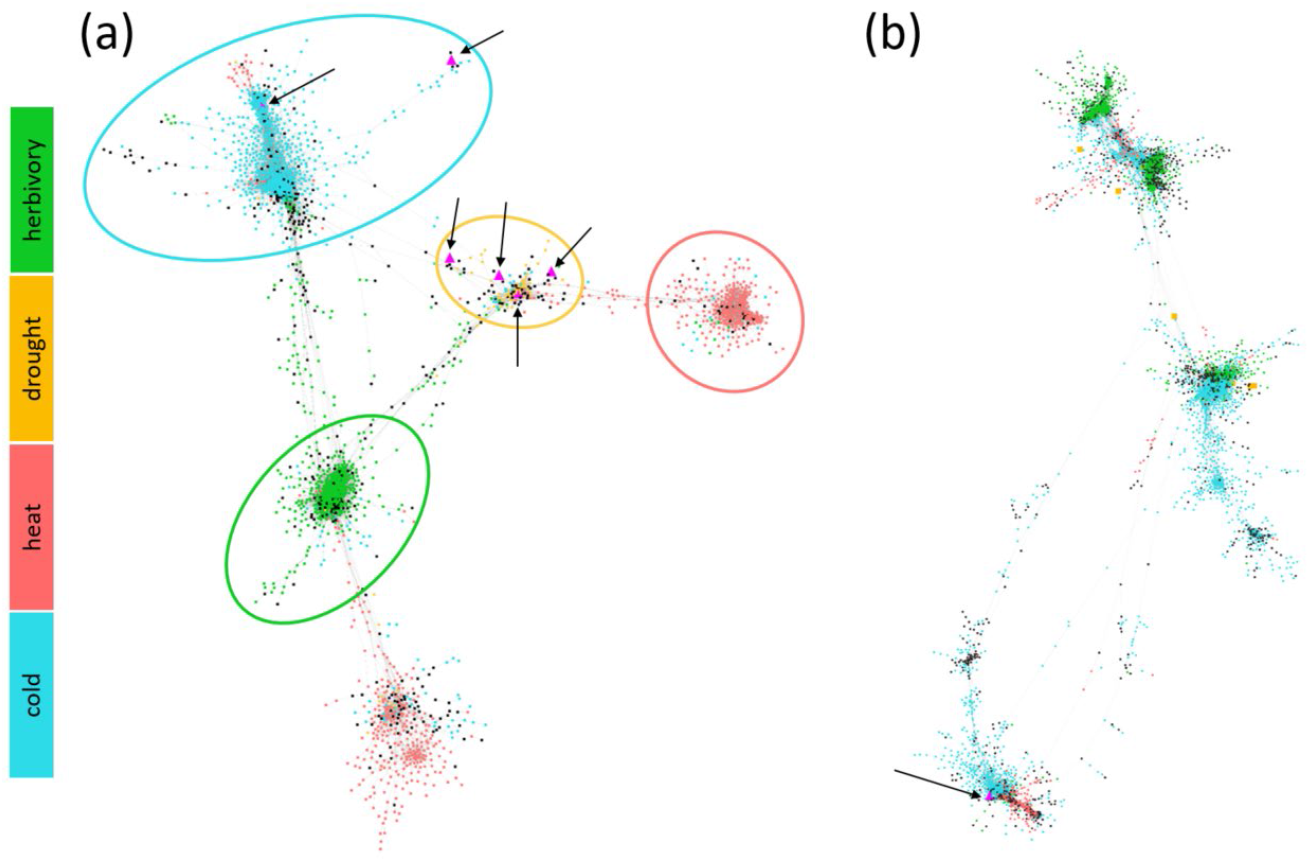
Co-expression network based on the transcriptional responses to cold, heat, drought and herbivory treatments (i.e. DEG-networks) in (a) *Biscutella laevigata* and (b) *Arabidopsis thaliana*. Differentially expressed genes (DEGs) that are strongly co-expressed across environmental treatments cluster according to the correlation of their expression profiles. Treatment-specific DEGs are coloured accordingly, whereas DEGs shared among treatments are displayed in black. Subnetworks in *B. laevigata* comprise of treatment-specific DEGs, while most down-regulated genes cluster in the non-circled subnetwork at the bottom. Magenta triangles (highlighted by arrows) mark the locations of DEGs that are up-regulated in response to all treatments and locate predominantly in the drought-subnetwork, indicative of drought responsive genes being equally involved in all treatments.

Most of the 1’504 DEGs identified in both *B. laevigata* and *A. thaliana* (i.e. 1’351 and 888 DEGs, respectively) appeared highly connected and were located across DEG-networks (SI figure 6). Out of the 66 environment-responsive transcription factors identified in both species, 15 and 31 were highly connected to their respective network and highlighted a conserved set of environment-responsive genes connected to species-specific DEGs.

Synergistic DEGs appeared predominantly located at the intersections of treatment-specific subnetworks in *B. laevigata*, highlighting their role in multiple transcriptional responses (figure 4a). The drought subnetwork comprised several such DEGs, as well as four of the seven genes up-regulated in all treatments, highlighting the synergistic role of drought-DEGs to environmental changes. This further contrasted with the few synergistic DEGs of *A. thaliana*, which appeared located among several loose clusters rather than at the intersections between clusters (figure 4b). Trade-off DEGs were contrastingly located within treatment specific subnetworks in *B. laevigata*, whereas they were distributed across the DEG-network in *A. thaliana*.

**Figure 4.**
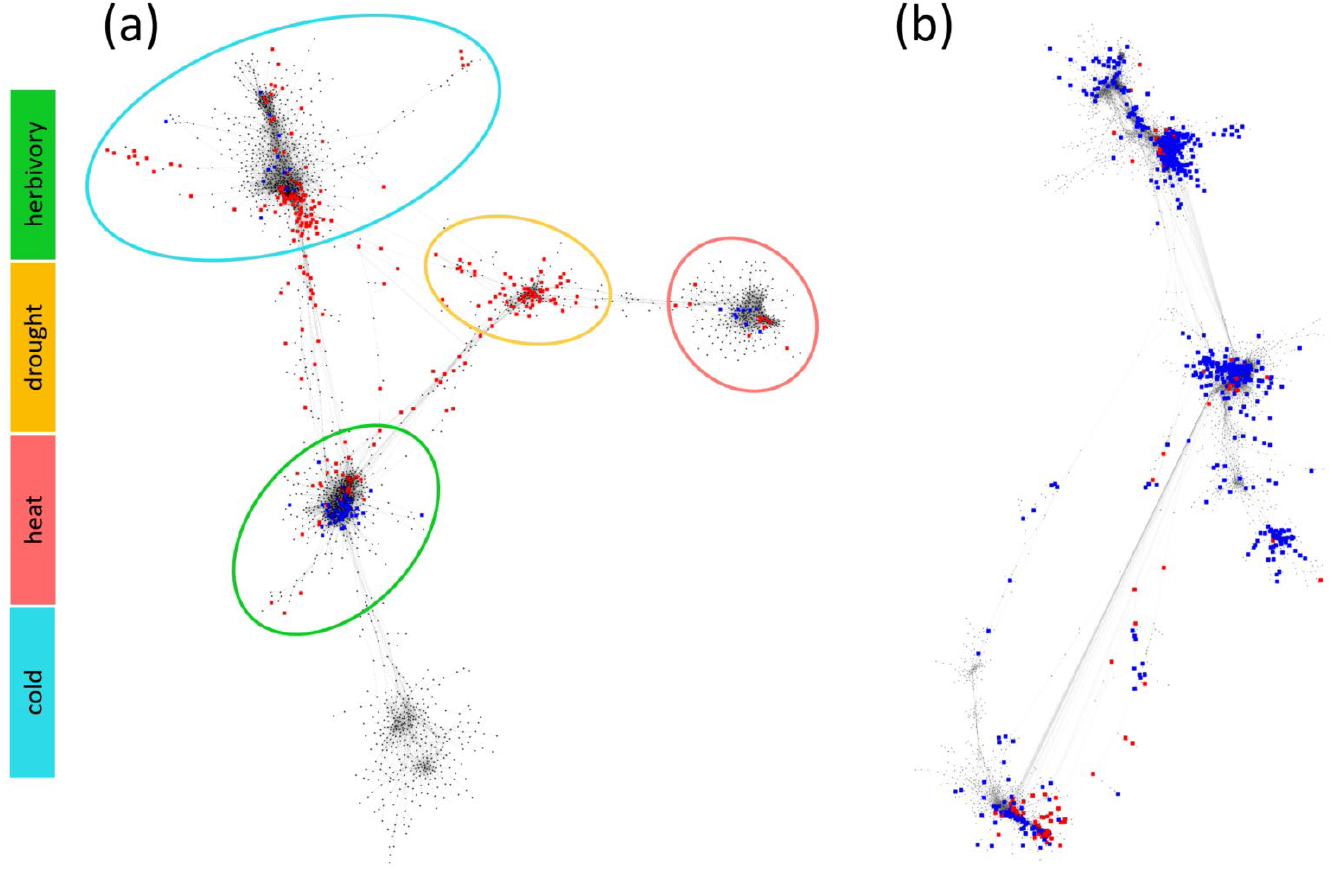
Architecture of the *Biscutella laevigata* and *Arabidopsis thaliana* DEG-networks. Synergistic DEGs (i.e. up-regulated in at least two treatments) are highlighted in red, whereas trade-off DEGs (i.e. up-regulated in one treatment and down-regulated in another) are highlighted in blue, showing the contrasted network architecture of the transcriptional responses to environmental treatments in *B. laevigata* and *A. thaliana*.**a)** The *B. laevigata* DEG-network showing 357 synergistic DEGs at the intersection of treatment-specific clusters and 105 trade-off DEGs locating within the cold, heat and herbivory but not the drought subnetwork. **b)** The *A. thaliana* DEG-network contrastingly presents only 285 synergistic DEGs, but 999 trade-off DEGs located across the entire network, indicative of considerably less synergistic transcriptional responses to investigated treatments.

## Discussion

Promoting accurate characterization of transcriptional changes in non-model organisms responding to environmental changes, RNAseq approaches offer crucial functional insights and foster our understanding of phenotypic plasticity. Relying on a replicated experimental design supporting reliable conclusions (Conesa *et al*., 2016), the quantification of expression based on both a *de novo* transcriptome or a genome assembly here highlighted similar transcriptional changes in response to cold, heat, drought and herbivory. As evaluated by publicly available data from the related model plant *A. thaliana* subjected to same environmental treatments, credible patterns of gene expression were identified and supported sensitive insights to be gathered by RNAseq in non-model organisms. However, focusing on loci expressed ≥1 TPM (12.3% of all *de novo* assembled genes) appeared necessary to evaluate biologically meaningful transcripts and promote comparability of *de novo*-vs genome-based insights. As signals of differential expression may be under-estimated with the former approach, mainly due to inaccurate resolution of gene models (Li & Dewey, 2011), genome-based quantification of gene expression should however be favoured when possible (Wang & Gribskov, 2017).

### Plant transcriptional responses to environmental changes

Gene expression was here quantified late after the onset of environmental changes and thus likely characterized new transcriptional steady states, potentially emphasizing on newly reached homeostasis rather than early signalling and responses (Kollist *et al*., 2019; Zhang *et al*., 2022). Accordingly, differential expression of typical early responding genes, such as *CBF1-3* under cold (Park *et al*., 2015) was detected in neither species, while downstream targets such as *STCH4* (Yu *et al*., 2020) were induced. Functions of the 1’504 DEGs shared by *B. laevigata* and *A. thaliana*, including 66 DNA-templated transcription factors, were indicative of changes underlying resource allocation and growth in response to environmental changes. In particular, the induction of circadian clock genes *CCA1* and *LHY* (Kyung *et al*., 2022) support a role of starch mobilisation to not only supply energy for growth under abiotic stresses (Moraes *et al*., 2022), but also as a source of osmolytes controlling stomata opening (Thalmann & Santelia, 2017). Accordingly, synergistic up-regulation of the starch degrading β-amylase *BAM1* under our cold and heat treatments, as well as up-regulation of *BAM3* under cold, indicate that both species rely on leaf starch during abiotic treatments tested here.

While herbivory treatments differed by the use of the generalist *P. xylostella* in *B. laevigata* and the Brassicaceae specialist *P. rapae* in *A. thaliana*, relatively few herbivore-specific transcriptional changes were detected, as expected by similar responses to *P. rapae* and the generalist *Spodoptera littoralis* in *A. thaliana* (Reymond *et al*., 2004). Several herbivory-responsive genes presented contrasted expression changes between species, with *P. xylostella* inducing up-regulation of homologues of *TPS03* (producing defense-related terpenoids; Huang *et al*., 2010; Knauer *et al*., 2018) in *B. laevigata*, whereas it was down-regulated following attack by *P. rapae* in *A. thaliana*. Similarly, as expected following the evolutionary arm race between specialist herbivores and Brassicaceae (Edger *et al*., 2015), the enzyme catalysing the first step in glucosinolate biosynthesis *CYP79B2* was also strongly up-regulated in *B. laevigata*, but un-induced by *P. rapae* in *A. thaliana*.

Differential expression of TEs in *B. laevigata* identified clades of interspersed *Copia* retrotransposons being up-regulated following a specific environmental treatment and indicating TE transcriptional activation (Grandbastien, 2015). Although only loosely related to the iconic heat-induced TE *ONSEN* (Ito *et al*., 2011), copies of *TERESTRA* were shown as massively up-regulated under heat in *B. laevigata*,as they also do in other Brassicaceae (Pietzenuk *et al*., 2016). Similarly, 18 copies of a Copia retrotransposon homologous to *Atcopia93* (or *EVD*) were identified as herbivory-induced, consistent with this TE being known as transcriptionally and transpositionally activated in *A. thaliana* (Marí-Ordóñez *et al*., 2013) following plant immune response (Zervudacki *et al*., 2018). Contrastingly, nine related copies of a *Copia* retrotransposon homologous to *Atcopia31* were identified as specifically herbivory-induced in *B. laevigata*, although it was surmised as heat-activated in *A. thaliana* (Quadrana *et al*., 2016; Pietzenuk *et al*., 2016). Although neither the extent to which such environment-responsive TE transcription supports transposition events, nor potential consequences of such regulation in natural population is known, RNAseq enabled an accurate characterization of responses of TEs to environmental changes in a non-model species.

### Modular transcriptional changes under environmental changes

The overall up-regulation of genes and the stronger shifts in expression in *B. laevigata* markedly contrasted with chiefly down-regulation and weaker shifts in *A. thaliana* towards new transcriptional steady states under environmental changes. Over 50% DEGs in *A. thaliana* showed constitutive expression in *B. laevigata*, suggesting that the stress-tolerant alpine species is constantly expressing genes used to cope with specific environmental changes in the model species. To what extent such constitutive expression is costly and adaptive, explaining the slow growth and limited competitive ability of *B. laevigata*, deserves additional work.

Although presenting the smallest number of DEGs in both species, the drought treatment involved unusually large proportions of DEGs that shifted their expression under at least another treatment, suggesting that genes related to water homeostasis are central to transcriptional responses to environmental changes. Consistent with a key role of drought-responsive genes in transcriptional responses to other abiotic and biotic stressors, they showed particularly strong expression shifts in *B. laevigata* and formed a central DEG-subnetwork involving a majority of the up-regulated DEGs by all treatments, including several other synergistic DEGs, but none of the identified trade-off DEGs.

Modules of co-expressed genes further characterized the architecture of transcriptional changes in response to cold, heat, drought and herbivory. In particular, the highly-structured DEG-network in *B. laevigata* presented clearly distinct subnetworks of co-regulated genes in response to each treatment and limited interactions among environment-specific subnetworks, supporting a modular transcriptional plasticity in response to environmental changes in *B. laevigata*. Although the extent to which such a highly modular transcriptional changes promote stress tolerance in *B. laevigata* remains out of scope, it sharply contrasts the observed architecture of transcriptional changes in *A. thaliana*, where up-DEGs responding to one environmental treatment were subnetwork-specific and synergistic DEGs were located within clusters rather than at their intersections. Consistent with highly integrated transcriptional responses to environmental changes, the fast-cycling *A. thaliana* further presented a considerably higher number of trade-off DEGs distributed across the entire network, compared to the few trade-off DEGs central to treatment-specific subnetworks in *B. laevigata*. As expected under decreasing herbivory pressure in colder climates (Rasmann *et al*., 2014), the largest proportion of trade-off DEGs were involved in unlikely simultaneous cold and herbivory stressors, in both species. Although experiments involving several unique treatments offer key transcriptional insights, multiple environmental stressors can be expected to simultaneously occur in nature. These result in non-additive transcriptional responses that remain to be considered, to fully understand transcriptional plasticity (Atkinson & Urwin, 2012; Suzuki *et al*., 2014; Prasch & Sonnewald, 2015).

As both species differ by multiple traits, underpinnings of their different transcriptional responses to environmental changes remain elusive. However, *B. laevigata* is a long-lived perennial that strategically allocates resources to tolerate stress while maintaining growth and reproduction, whereas the short-lived *A. thaliana* likely maximizes growth and reproduction by avoiding environmental stresses (Lundgren & des Marais, 2020). To what extent different ecological strategies (Grime, 1977; Dìaz *et al*., 2016) and costs associated with plastic stress tolerance select for different transcriptional architectures shall be addresses with similar data from a larger set of species. Furthermore, *B. laevigata* has undergone an additional whole-genome duplication event as compared to *A. thaliana* (Geiser *et al*., 2016) and numerous environment-responsive genes were here found with multiple homologues showing variable responses to treatments. To what extent the initial redundancy resulting from gene duplication supports robustness (Wagner, 1994, 2002) and contributed to the evolution of modular transcriptional responses remains to be investigated.

### Data statement

The 17 RNAseq raw read files of the *B. laevigata* leaf transcriptomes under environmental stress are available in the European Nucleotide Archive (ENA accession: PRJEB48469). The seven RNAseq raw read files of the *B. laevigata* tissue atlas are available under ENA accession PRJEB48599.

## Acknowledgements

We thank C. Ball and J. Sekulovski for taking excellent care of our plants, C. Robert for support in designing the herbivory experiment and T. Züst for support in data analysis. Thanks to V. Ernst, T. Bürki, V. Pulver, A. Metry for advice and fruitful discussions. Thanks to S. Grünig, N. Schenk and J. Schröder for helpful comments on the manuscript. This research was funded by the Swiss National Science Foundation (Grant 31003A_178938).

## Author contributions

MB and CP designed the research; MB and RRC collected data, MB and BM analysed data, MB and CP interpreted data; MB and CP wrote the manuscript.

## Supporting Information

SI tables 1 and 2: List of *B. laevigata* and *A. thaliana* RNAseq libraries.

SI tables 3 and 4: Mapping rates of the 17 *B. laevigata* environmental treatment RNAseq libraries to the *de novo* transcriptome and genome assembly.

SI tables 5: Mapping rates of the 19 *A. thaliana* RNAseq libraries to the Araport11 coding sequences.

SI tables 6 and 7: Lists of *B. laevigata* and *A. thaliana* DEGs with log2-fold-changes.

SI table 8: Proximity of genes to environmentally induced transposable elements.

SI tables 9-25: GO-enrichment of DEGs under cold, heat, drought and herbivory.

SI table 26: Annotation of the transcription factors shared between *B. laevigata* and *A. thaliana*.

SI tables 27-33: GO-enrichment of synergistic DEGs, trade-off DEGs, species specific DEGs and *A. thaliana* DEGs with constitutive expression in *B. laevigata*.

SI tables 34 and 35: Co-expression modules and their correlations to treatments.

SI figure 1. Number of expressed genes in tissues used for the *Biscutella laevigata de novo* transcriptome assembly.

SI figure 2. Setup of the herbivory treatment for *Biscutella laevigata* and feeding damage caused by *Plutella xylostella* larvae.

SI figure 3. Absolute log2-fold changes (logFC) of differentially expressed genes in *Biscutella leavigata* and *Arabidopsis thaliana* under environmental treatments (cold, heat, drought and herbivory).

SI figure 4. The *Biscutella laevigata* co-expression network with differentially expressed genes in response to cold, heat, drought and herbivory treatments forming separate clusters.

SI figure 5. The *Arabidopsis thaliana* co-expression network with differentially expressed genes in cold, heat, drought and herbivory treatments highlighted in red.

SI figure 6. Differentially expressed genes shared by *Biscutella laevigata* and *Arabidopsis thaliana* highlighted across co-expression networks.

Supplementary text T1: Central synergistic DEGs and their network neighbours

