## Supplementary Information for "Modular transcriptional responses to environmental changes"

***SI table of contents:***

SI tables 1 and 2: List of *B. laevigata* and *A. thaliana* RNAseq libraries.

SI tables 3 and 4: Mapping rates of the 17 *B. laevigata* environmental treatment RNAseq libraries to the *de novo* transcriptome and genome assembly.

SI tables 5: Mapping rates of the 19 *A. thaliana* RNAseq libraries to the Araport11 coding sequences.

SI tables 6 and 7: Lists of *B. laevigata* and *A. thaliana* DEGs with log_2_-fold-changes.

SI table 8: Proximity of genes to environmentally induced transposable elements.

SI tables 9-25: GO-enrichment of DEGs under cold, heat, drought and herbivory.

SI table 26: Annotation of the transcription factors shared between *B. laevigata* and *A. thaliana*.

SI tables 27-33: GO-enrichment of synergistic DEGs, trade-off DEGs, species specific DEGs and *A. thaliana* DEGs with constitutive expression in *B. laevigata*.

SI tables 34 and 35: Co-expression modules and their correlations to treatments.

SI figure 1. Number of expressed genes in tissues used for the *Biscutella laevigata* *de novo* transcriptome assembly.

SI figure 2. Setup of the herbivory treatment for *Biscutella laevigata* and feeding damage caused by *Plutella xylostella* larvae.

SI figure 3. Absolute log_2_-fold changes (logFC) of differentially expressed genes in *Biscutella leavigata* and *Arabidopsis thaliana* under environmental treatments (cold, heat, drought and herbivory).

SI figure 4. The *Biscutella laevigata* co-expression network with differentially expressed genes in response to cold, heat, drought and herbivory treatments forming separate clusters.

SI figure 5. The *Arabidopsis thaliana* co-expression network with differentially expressed genes in cold, heat, drought and herbivory treatments highlighted in red.

SI figure 6. Differentially expressed genes shared by *Biscutella laevigata* and *Arabidopsis thaliana* highlighted across co-expression networks.

Supplementary text T1: Central synergistic DEGs and their network neighbours

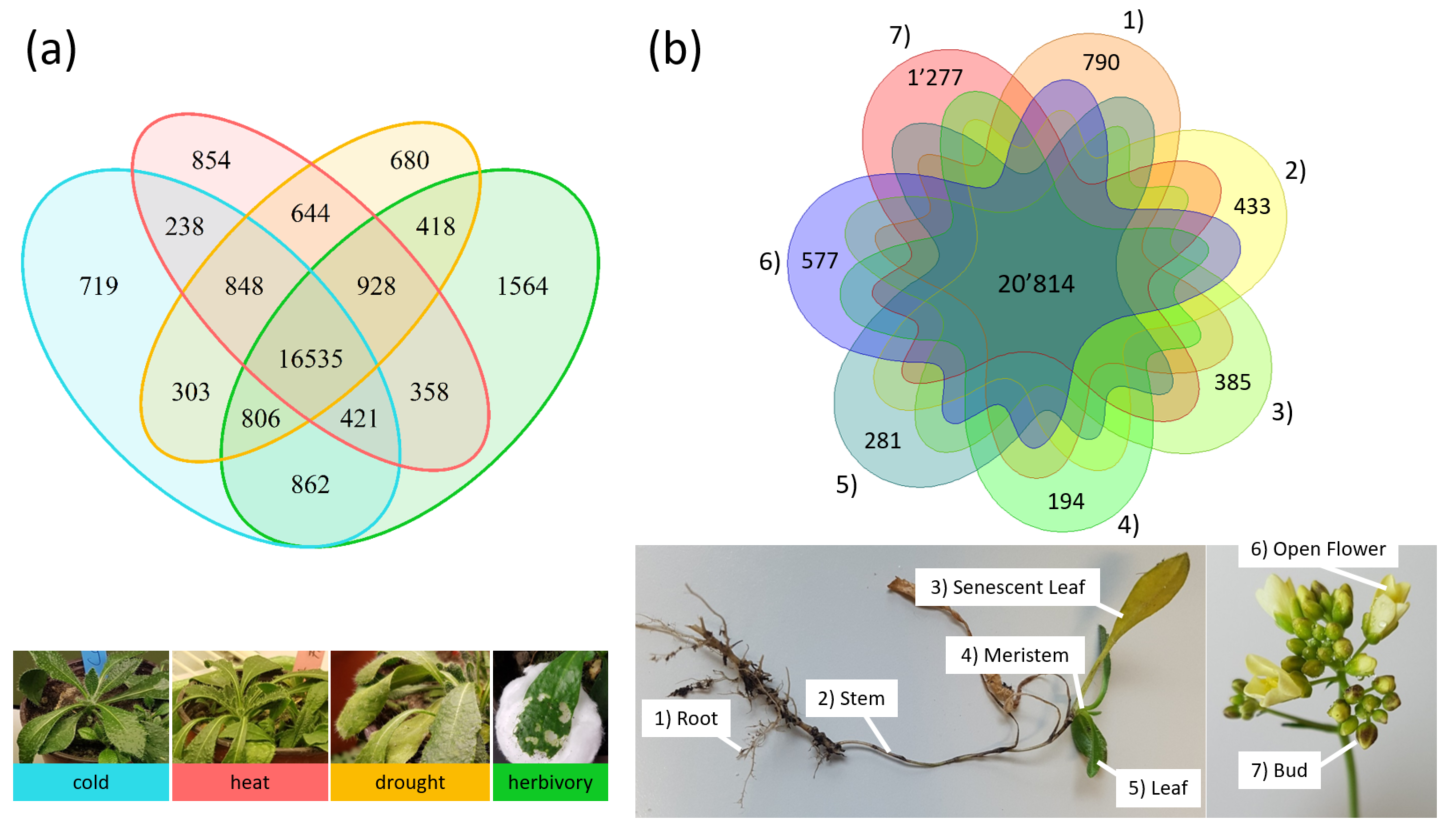

**SI figure 1. Number of expressed genes in tissues used for the *Biscutella laevigata de novo* transcriptome assembly.** The *B. laevigata de novo* transcriptome assembly was built from sequenced RNA reads of seven different tissues (root, stem, senescent leaf, meristem, leaf, open flower and bud) under control treatment as well as the library with the most reads from leaves under each environmental treatment (control, cold, heat, drought, herbivory). **a)** Venn diagram showing the number of genes (of 54’457 in the *B. laevigata* genome) expressed on average ≥1TPM in each treatment; cold: 20’732 (38.1%), heat: 20’826 (38.2%), drought: 21’162 (38.9%), herbivory : 21’892 (40.2%). **b)** Venn diagram showing the number of genes expressed ≥1TPM in each of the seven tissues; Root = 29’290 (53.8%); Stem = 27’874 (51.2%); Senescent Leaf = 26’979 (49.5%); Meristem = 29’057 (53.4%); Leaf = 27’448 (50.4%); Open Flower = 28’670 (52.6%) and Bud = 30’434 (55.9%). A total of 20’814 (38.2%) genes were commonly expressed in all tissues and between 194 to 1’277 (0.4-2.3%) genes were expressed specific to tissues.

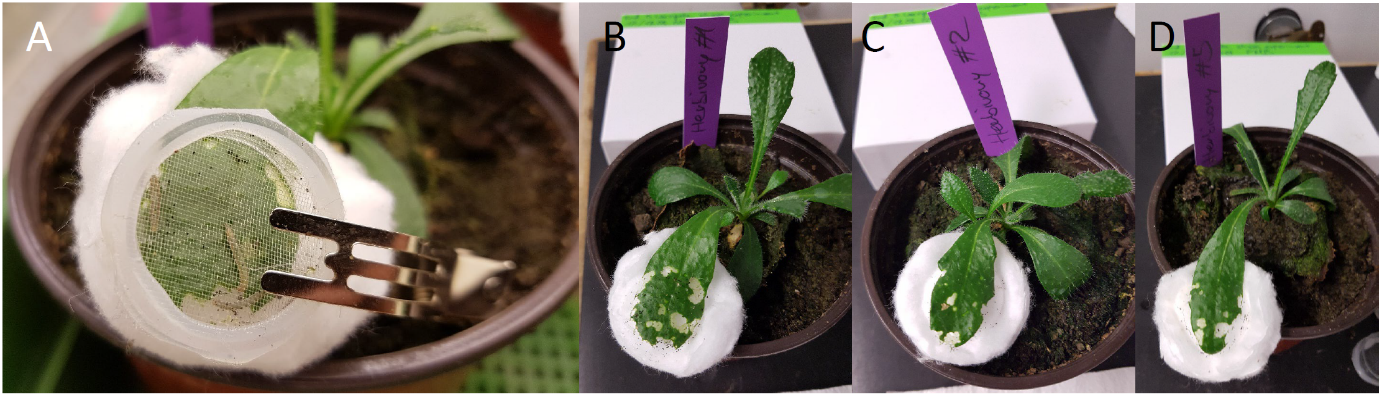

**SI figure 2. Setup of the herbivory treatment for *Biscutella laevigata* and feeding damage caused by *Plutella xylostella* larvae. a)** Small plastic cages were used to hold eight 3^rd^-5^th^ instar larvae of *P. xylostella* during the herbivory treatment on one leaf of *B. laevigata*. **b-d)** Feeding damage caused after 30h of herbivory on leaves used for RNA extraction and subsequent library preparation.

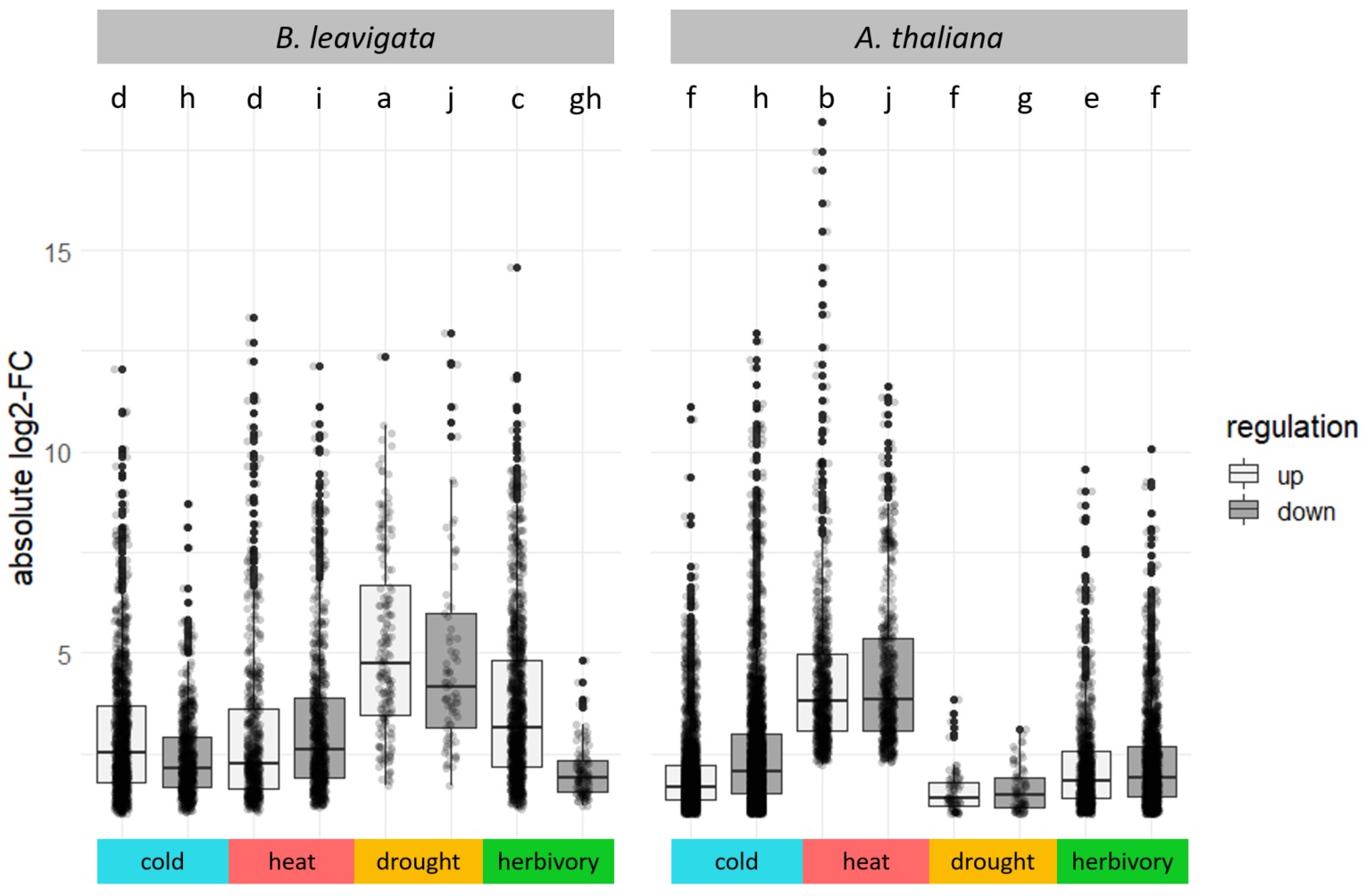

**SI figure 3. Absolute log_2_-fold changes (logFC) of differentially expressed genes in *Biscutella leavigata* and *Arabidopsis thaliana* under environmental treatments (cold, heat, drought and herbivory).** In *B. laevigata*, drought induces the strongest fold-changes, while in *A. thaliana* the strongest changes in gene expression are induced by the heat treatment. Analysis of variance (ANOVA) with a posthoc Tukey’s test on logFC values identified significantly different groups here shown as not sharing a letter on the top.

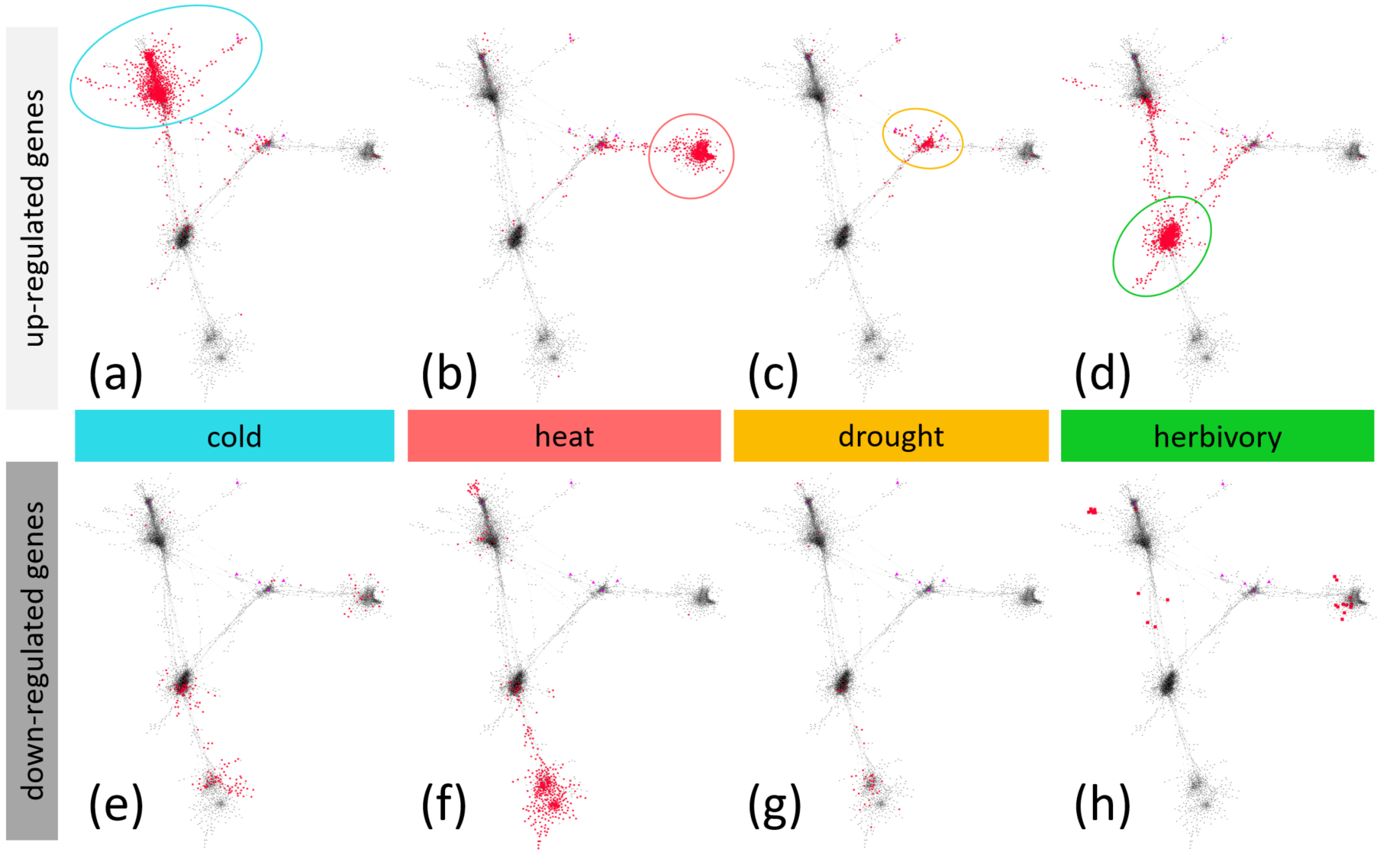

**SI figure 4. The *Biscutella laevigata* DEG-network with differentially expressed genes in response to cold, heat, drought and herbivory treatments forming separate clusters.** From the co-expression network consisting of 41’613 genes, 2’625 DEGs with an edge-weight ≥0.5 were extracted and visualized in Cytoscape. Up-regulated DEGs in response to **(a)** cold, **(b)** heat, **(c)** drought and **(d)** herbivory are highlighted in red and form four clusters (coloured circles). Cold, heat and herbivory up-regulated genes are all connected to the cluster of drought-responsive genes, highlighting their central role in all investigated treatments. Herbivory triggers the up-regulation **(d)** of many genes involved in the cold and drought responses. Down-regulated DEGs in response to **(e)** cold, **(f)** heat, **(g)** and drought are highlighted in red and mostly cluster together in the bottom subnetwork, while the few herbivory down-regulated DEGs **(h)** are mostly located in the cold subnetwork **(a)** and the heat subnetwork **(d)**. Down-regulated genes under cold **(e)**, heat **(f)** and drought **(g)** are to some extent located in the subnetworks of up-regulated genes, indicating that those genes are involved in expression trade-offs between treatments.

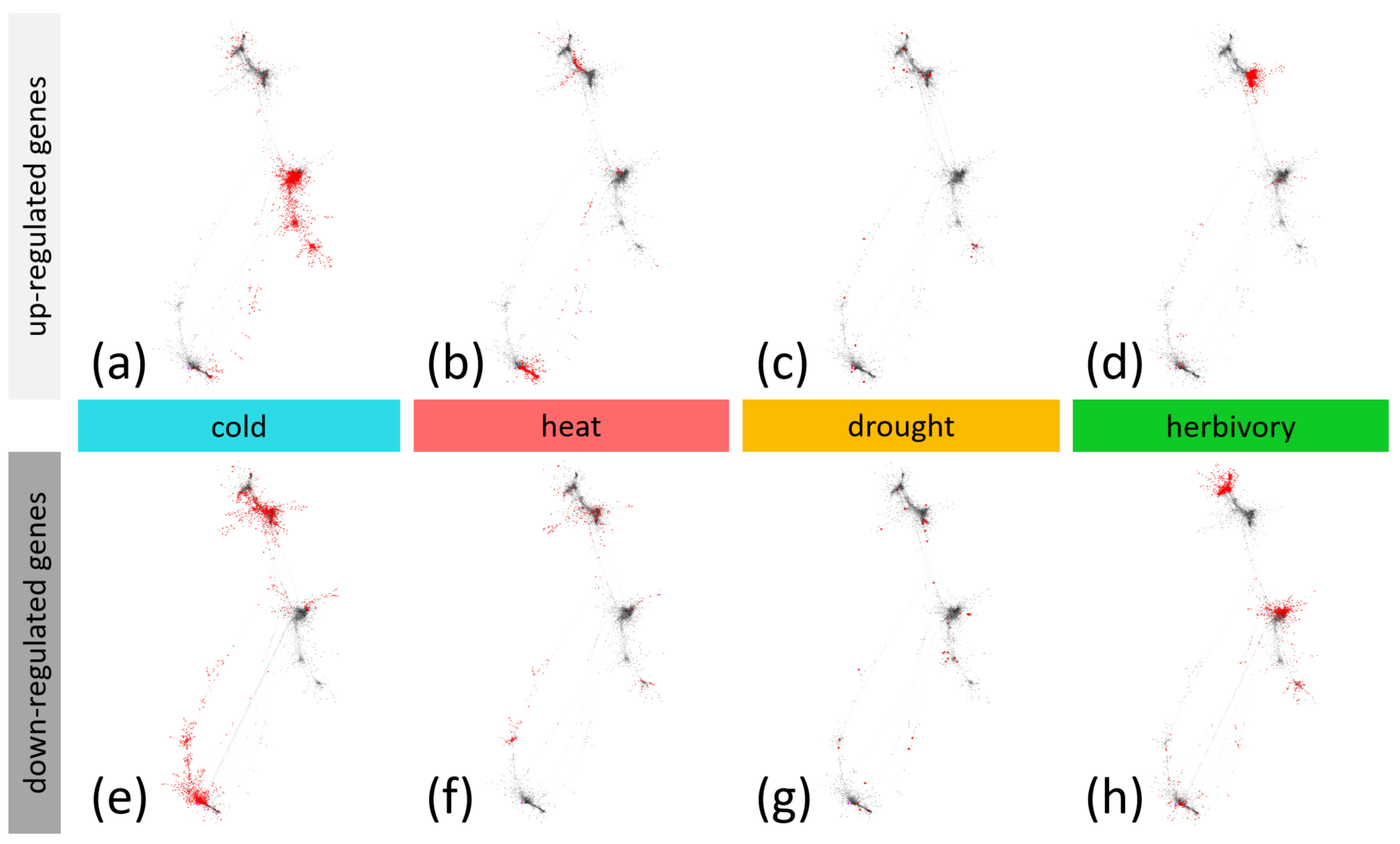

**SI figure 5**. **The *Arabidopsis thaliana* DEG-network with differentially expressed genes in cold, heat, drought and herbivory treatments highlighted in red.** From the *A. thaliana* co-expression network consisting of 26’051 genes, 4’465 DEGs with connections stronger than an edge-weight of 0.7 were extracted and visualized in Cytoscape. Genes do not cluster well according to the treatment they are up-regulated in. **a-d)** Up-regulated DEGs highlighted in red. **e-h)** Down-regulated DEGs highlighted in red.

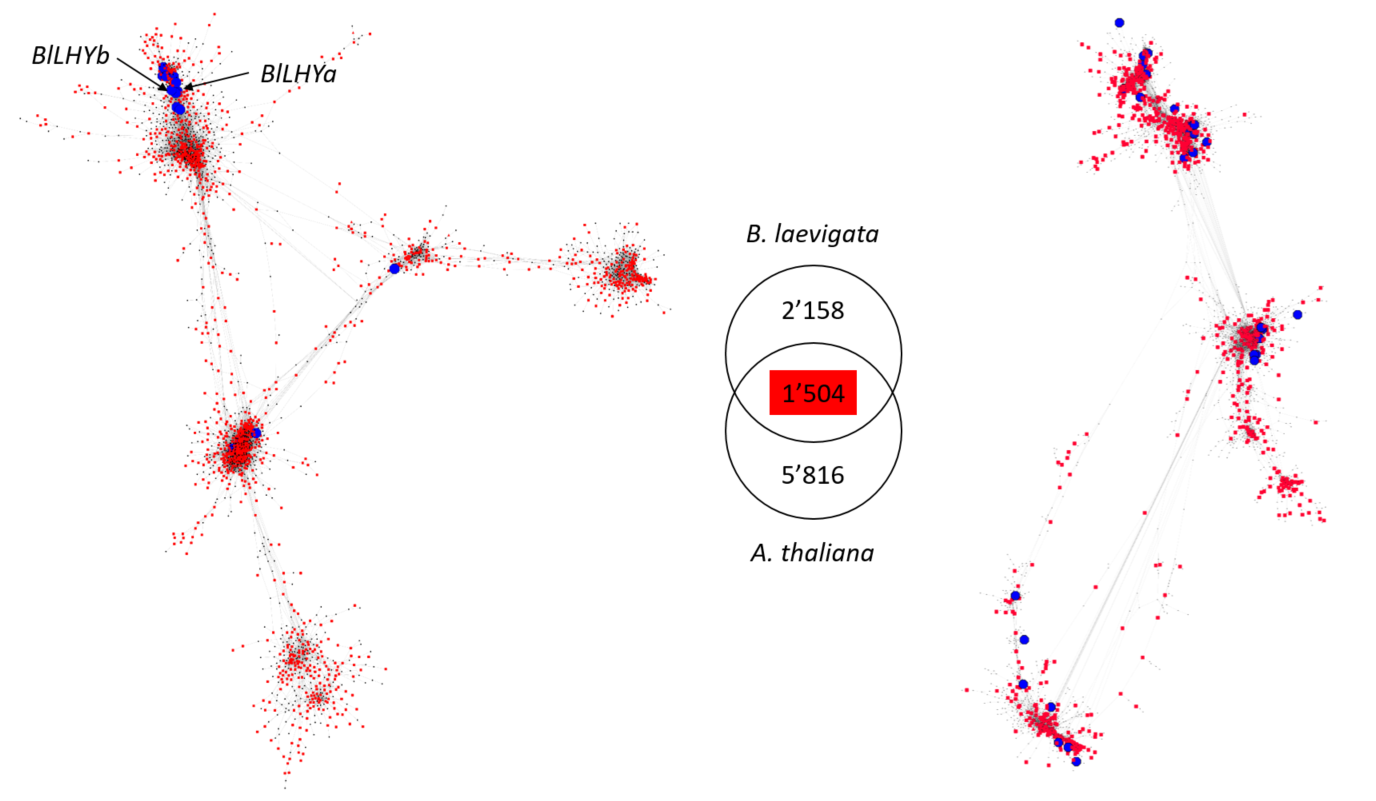

**SI Figure 6. Differentially expressed genes (DEGs) shared by *Biscutella laevigata* and *Arabidopsis thaliana* highlighted across co-expression networks.** The 1’504 DEGs shared between *B. laevigata* and *A. thaliana* (highlighted in red) were relatively even distributed across the DEG-networks. 1’351 and 888 of those shared DEGs were present in the *B. laevigata* and *A. thaliana* DEG-networks respectively. The shared DEGs contained 66 transcription factors (TFs), 15 and 31 of which were present in the *B. laevigata* and *A. thaliana* DEG-networks respectively (highlighted in blue). For example, we found two *LHY* homologues (*BlLHYa* and *BlLHYb*) located in the *B. laevigata* cold subnetwork. DEGs specific to either species are represented in black nodes in the respective DEG-networks. Noticeably, the 1’504 shared DEGs are represented by 2’014 homologues in *B. laevigata*, which is why the numbers of the Venn-diagram don’t add up to the total 4’172 DEGs of *B. laevigata*.

***Supplementary Text 1:* Central synergistic DEGs and their network neighbours**

Seven genes were highlighted as up-regulated in parallel in response to all environmental treatments in *B. laevigata* (SI figure T1). We characterized these stress-common-DEGs based on homology with *A. thaliana*, their clustering into co-expression modules identified in *B. laevigata*, as well as the functions of their direct neighbours in the co-expression network. As detailed below, all seven genes were involved in water homeostasis and response to abscisic acid and four of them were located in the central drought subnetwork. Contrastingly, *A. thaliana* presented only one DEG that was up-regulated in parallel under all environmental treatments (i.e. the methionine gamma lyase *AtMGL*; AT1G64660).

For BLAEV0010256 we found no homologous gene sequences with a nucleotide BLAST and an e-value cut-off of 1e^-10^. We therefore blasted the coding sequence of this gene manually with a discontiguous megablast (NCBI BLAST) and found 76% sequence identity to *A. thaliana* *CAP160* (AT4G25580) (SI figure T1a-1). In the *B. laevigata* co-expression network, this gene was grouped into module 56 (SI table 34) which correlates at 0.71 with cold, as well as weakly negative with drought (cor = -0.27) and heat (cor = -0.29). The gene itself correlated 0.77 with cold. In the *B. laevigata* subnetwork, it was found in the center of the cold cluster, with 56 first order neighbours, showing high expression similarity to these direct neighbours (figure T1b-1). The functions of direct neighbours highlighted broad functional similarity of neighbours of BLAEV0010256 that were enriched in “regulation of transcription, DNA-templated”, “response to cold”, “response to wounding” and “response to water deprivation” as well as response to light, such as “response to UV-B”, “response to red light” or “circadian rhythm” (SI figure T2).

BLAEV0008757 (figure T1a-2) is homologous to *LEA14* (AT1G01470). LEA-proteins are known to be induced by ABA, cold- and drought-stress and confer desiccation and freezing tolerance (Ingram & Bartels, 1996; Thomashow, 1999). In the *B. laevigata* co-expression network, it is grouped into the module 36 (SI table 34), which correlates 0.55 with drought. The gene itself correlates 0.56 with cold. In the *B. laevigata* DEG-subnetwork it is located at the periphery of the cold cluster and has three first order neighbours (figure T1b-2). The first order neighbours were BLAEV0044594 (edge-weight = 0.64; no homology found), BLAEV0022187 (edge-weight = 0.64; homologous to AT1G78070, a Transducin/WD40 repeat-like superfamily protein involved in the osmotic stress response and BLAEV0014038 (edge-weight = 0.5; homologous to AT2G17840, *ERD7* shown to affect cell membrane stability during cold stress (Barajas-Lopez *et al.*, 2021).

BLAEV0041356 (figure T1a-3) is homologous to *MYB112* (AT1G48000) and *Oryza sativa MYB2*, a R2R3-MYB transcription factor, responsive to ABA whose over-expression supported higher salt, cold and dehydration tolerance as well as high sensitivity to ABA in rice (Yang *et al.*, 2012). The transcription factor *AtMYB112*/*OsMYB2* has been shown to activate genes involved in anthocyanin production during salinity and high light stress (Lotkowska *et al.*, 2015). In the weighted gene co-expression network of *B. laevigata*, it grouped into the module 24 correlating at 0.58 with cold (SI table 34). The gene itself did not significantly correlate with any treatment. It appeared strongly connected (edge-weights between 0.5 and 0.68) to four first order neighbours (figure T1b-3). The first order neighbours BLAEV0019155 and BLAEV0019156 are both homologous to AT5G05850, *PIRL1* intracellular Leucine rich repeat protein, shown to be essential for differentiation of microspores into pollen and hypothesized to play a role in signal transduction (Forsthoefel *et al.*, 2005, 2010). The gene BLAEV0050078 is homologous to AT3G61160, a GS3-like kinase known for a role in signal transduction during environmental stress and development (Richard *et al.*, 2005; Li *et al.*, 2021). The gene BLAEV0016556 is homologous to AT5G57910 or *INO80-BINDING PROTEIN 2B*, a ribosomal RNA small subunit methyltransferase-G binding to the INO80 chromatin remodelling complex (Shang *et al.*, 2021).

BLAEV0011935 (figure T1a-4) is homologous to *LEA46*. Found in module 36, which correlates 0.55 with drought (SI table 34). It has only one first order neighbour BLAEV0022112 (edge-weight = 0.57; homologous to AT1G77450, *ANAC032*) and is located at the center of the drought subnetwork. *ANAC032* is a transcription factor induced by ABA, oxidative stress and leaf senescence (Takasaki *et al.*, 2015; Mahmood *et al.*, 2016a,b). It inhibits photosynthesis, leads to accumulation of reactive oxygen species and carbon starvation (Sun *et al.*, 2019) and regulates stress responding genes in roots through activation of *MYB30* (Maki *et al.*, 2019).

BLAEV0051171 (figure T1a-5) is homologous to *ESL1* (AT1G08920), a sugar transporter of the “early response to dehydration six-like” gene family, that is commonly expressed under drought stress (Slawinski *et al.*, 2021). Found in module 5, which correlates 0.76 with drought (SI table 34), the gene itself correlates at 0.69 with drought. It is located at the center of the drought subnetwork in *B. laevigata*, with 53 first order neighbours (among which the above-mentioned *ANAC032*). Those 53 neighbours were enriched in the biological processes such as “response to water deprivation”, “response to abscisic acid” and “response to salt stress” (SI figure T3).

BLAEV0020146 (figure T1a-6) is homologous to AT5G15190, a gene coding for a hypothetical protein involved in the ABA-activated signalling pathway, which is annotated with GO-terms “response to water deprivation” and “response to wounding”. In the *B. laevigata* co-expression network, it is grouped into module 5 (SI table 34) and therefore shows co-expression with the above-mentioned BLAEV0051171 (*ESL1*) and its neighbours. The gene BLAEV0020146 correlates 0.52 with drought and is located in the periphery of the drought-subnetwork strongly connected to the BLAEV0020145 (edge weight = 0.73; unannotated) and BLAEV0005239 (edge weight = 0.61; homologous to AT2G39050, *EULS3* a lectin protein binding to carbohydrates in response to stress, particularly involved in stomatal closure (van Hove *et al.*, 2014, 2015).

BLAEV0001449 (figure T1c) is homologous to *PP2C49* (AT3G62260), a protein phosphatase known for its negative regulation of stress-induced MAP-kinase pathways (Schweighofer *et al.*, 2004). It is grouped into the module 36 (SI table 34) with the two above-mentioned homologues of *LEA14* (BLAEV0008757) and *LEA46* (BLAEV0011935). It is not strongly correlated with any of the environmental treatments tested here and is not strongly enough connected to any of the environmental response genes to be included in the *B. laevigata* co-expression network (edge-weight <0.5).

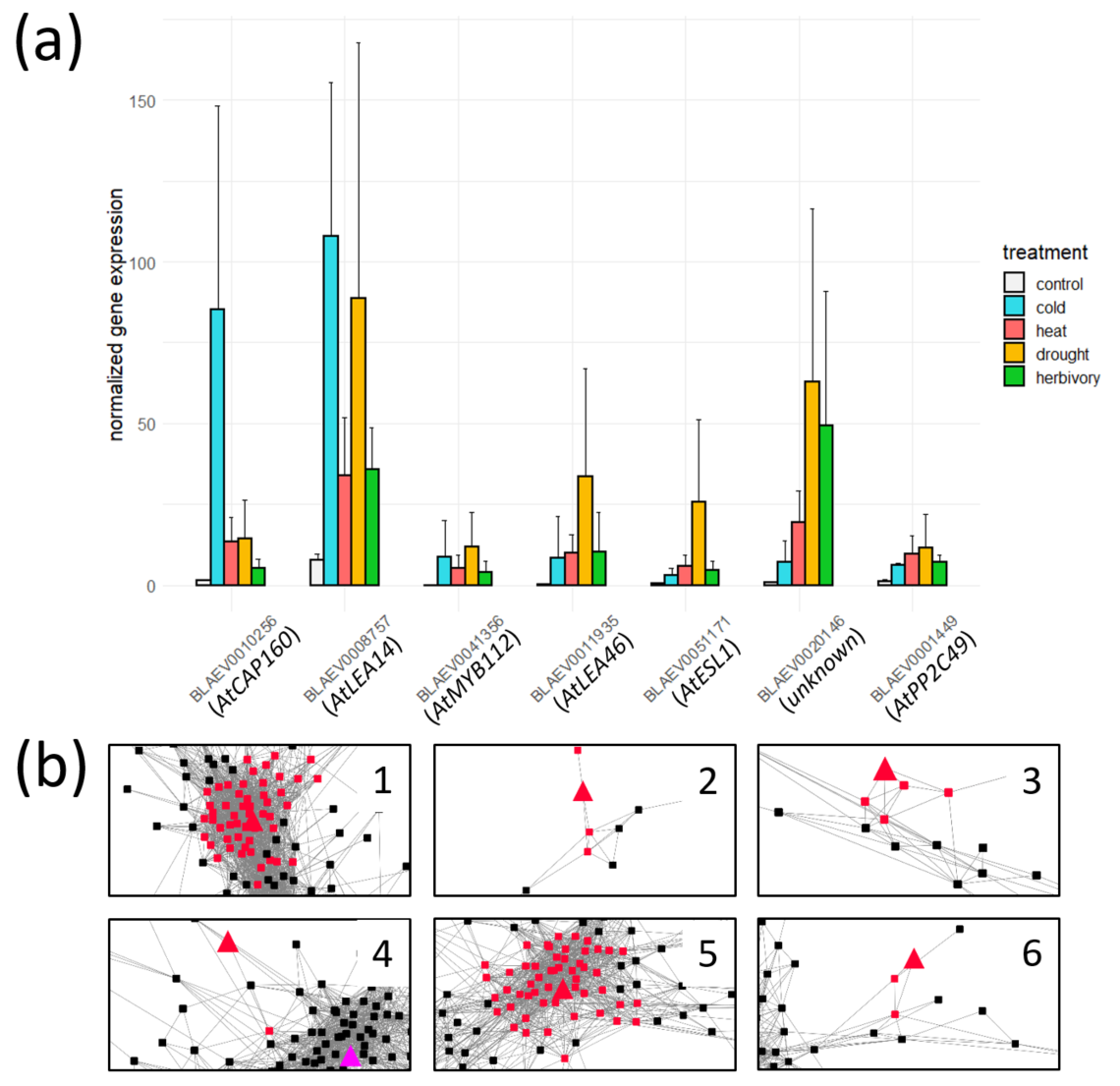

**SI figure T1. Differentially expressed genes (DEGs) commonly up-regulated in response to cold, heat, drought and herbivory in *Biscutella laevigata*. a)** Barplot showing normalized gene expression (TMM normalized TPM) for each of the seven DEGs up-regulated in each treatment. Labels refer to homologous genes in *A. thaliana*. The seventh gene *AtPP2C49* (BLAEV0001449) shows connection to other DEGs below 0.5 and is not included in the DEG-network. Error bars show SD. **b)** The *B. laevigata* DEG-network contains 6 of the 7 DEGs that are up-regulated in all stress treatments (shown as triangles). Four of these genes (3-6) locate in the drought DEG-subnetwork (see also figure 3). Direct neighbours of each of these six stress-common-DEGs are highlighted in red. **b1)** BLAEV0010256 is homologous to *AtCAP160* and connects to 56 direct neighbours (see also figure T2). **b2)** BLAEV0008757 is homologous to *AtLEA14*, 3 direct neighbours. **b3)** BLAEV0041356 is homologous to *AtMYB112*, 4 direct neighbours. **b4)** BLAEV0011935 is homologous to *AtLEA46*, 1 direct neighbour. **b5)** BLAEV0051171 is homologous to *At*ESL1, 53 direct neighbours (see also figure T3). **b6)** BLAEV0020146 is homologous to AT5G15190, a gene coding for a hypothetical protein involved in the ABA-activated signalling pathway, 2 direct neighbours.

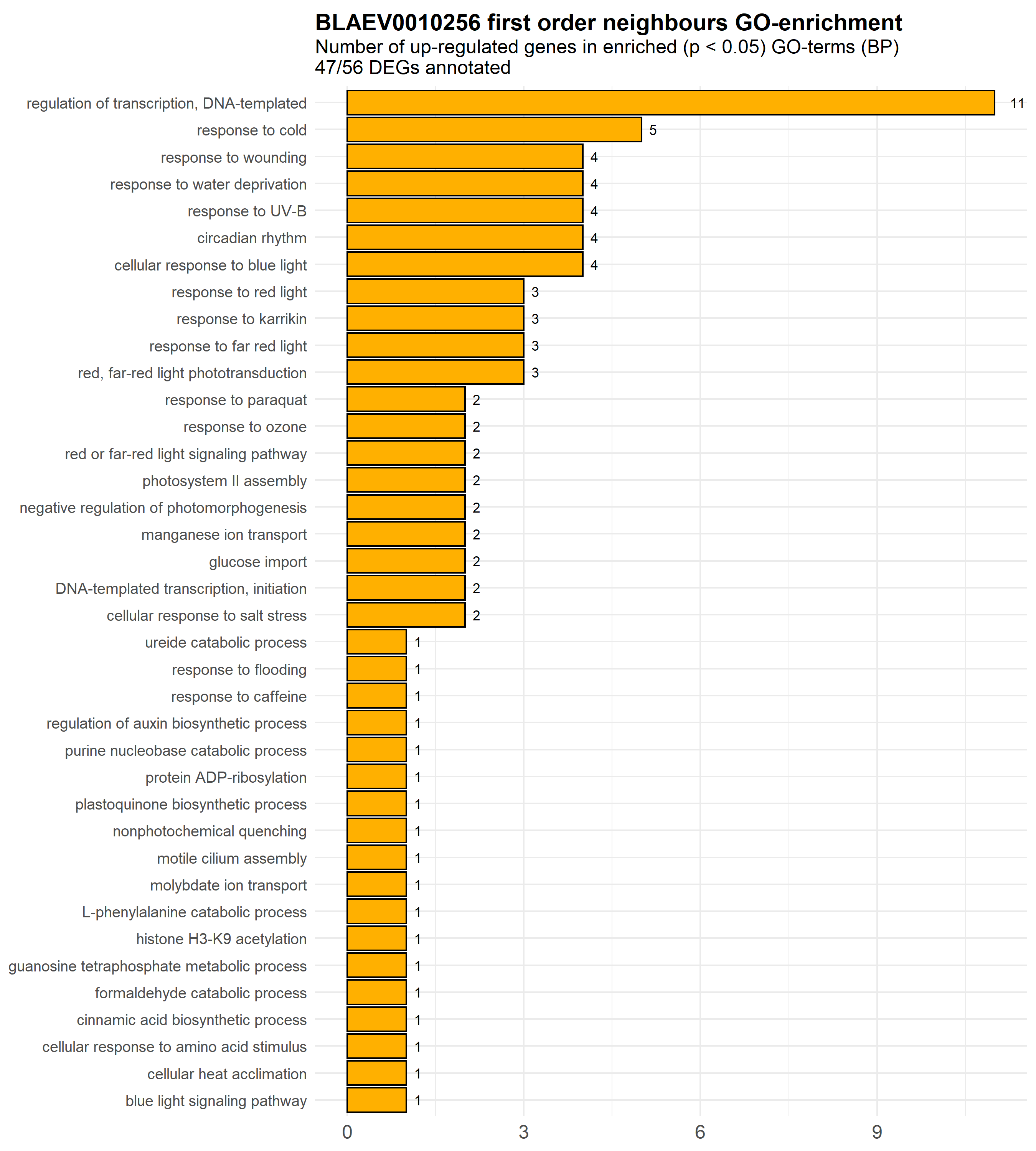
**SI figure T2. Gene Ontology (GO) enrichment results for the 56 direct neighbours of BLAEV0010256.** Direct neighbours of BLAEV0010256 are mostly DNA-templated transcription factors involved in the response to cold, drought and light conditions. GO-terms enriched at p-value <0.05 are depicted with the number of cold up-regulated genes that are annotated with each GO-term.

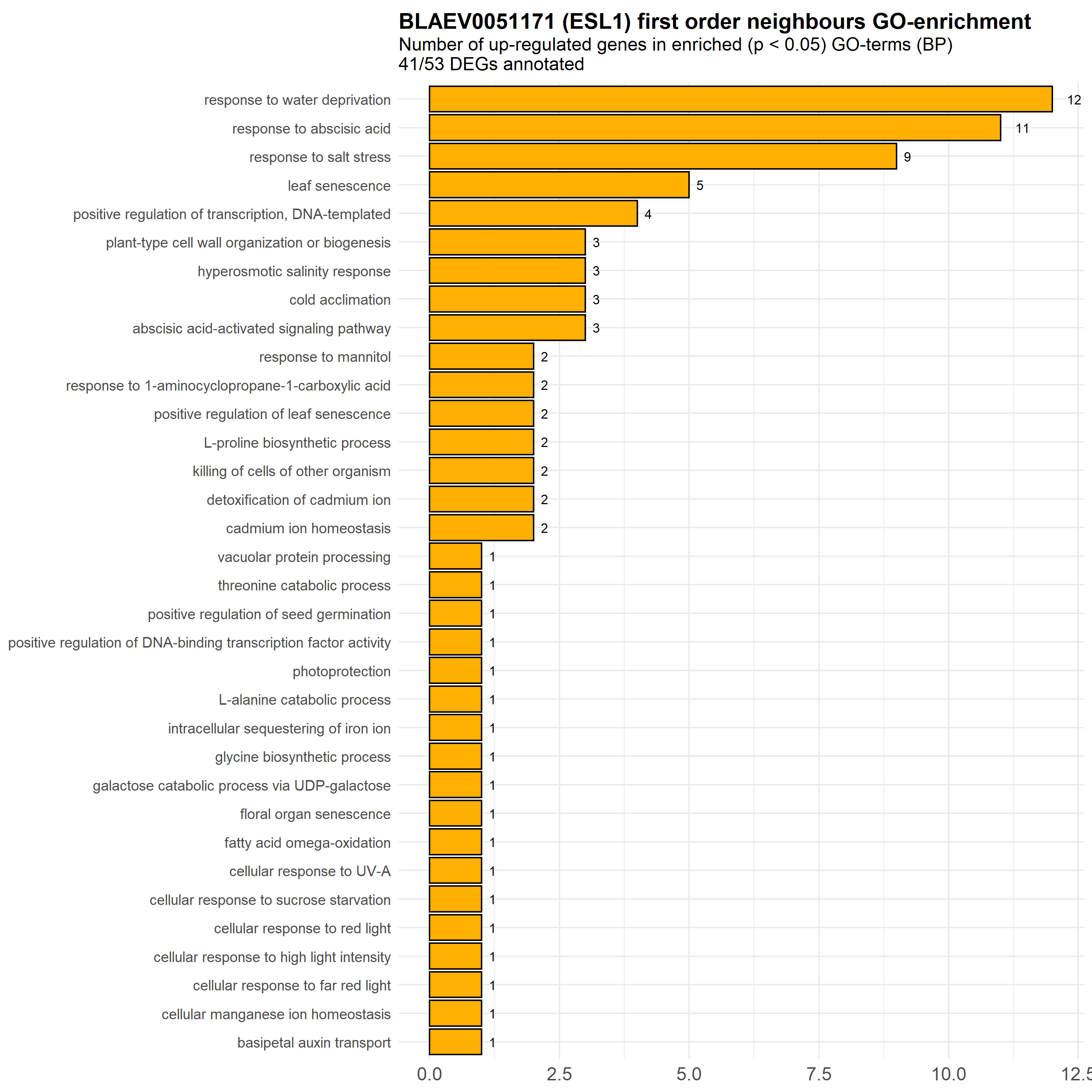
**SI figure T3. Gene Ontology (GO) enrichment results for the 53 direct neighbours of BLAEV0051171 (*ESL1*).** Direct neighbours of BLAEV0051171 are mostly involved the response to drought, as indicated by the response to water deprivation, salt stress and abscisic acid. GO-terms enriched at p-value <0.05 are depicted with the number of cold up-regulated genes that are annotated with each GO-term.

***SI references***

**Barajas-Lopez J de D, Tiwari A, Zarza X, Shaw MW, Pascual J, Punkkinen M, Bakowska JC, Munnik T, Fujii H**. **2021**. EARLY RESPONSE to DEHYDRATION 7 remodels cell membrane lipid composition during cold stress in Arabidopsis. *Plant and Cell Physiology* **62**: 80–91.

**Forsthoefel NR, Cutler K, Port MD, Yamamoto T, Vernon DM**. **2005**. PIRLs: A novel class of plant intracellular leucine-rich repeat proteins. *Plant and Cell Physiology* **46**: 913–922.

**Forsthoefel NR, Dao TP, Vernon DM**. **2010**. PIRL1 and PIRL9, encoding members of a novel plant-specific family of leucine-rich repeat proteins, are essential for differentiation of microspores into pollen. *Planta* **232**: 1101–1114.

**van Hove J, de Jaeger G, de Winne N, Guisez Y, van Damme EJM**. **2015**. The Arabidopsis lectin EULS3 is involved in stomatal closure. *Plant Science* **238**: 312–322.

**van Hove J, Stefanowicz K, de Schutter K, Eggermont L, Lannoo N, al Atalah B, van Damme EJM**. **2014**. Transcriptional profiling of the lectin ArathEULS3 from Arabidopsis thaliana toward abiotic stresses. *Journal of Plant Physiology* **171**: 1763–1773.

**Ingram J, Bartels D**. **1996**. The molecular basis of dehydration tolerance in plants. *Annu. Rev. Plant Physiol. Plant Mol. Biol* **47**: 377–403.

**Lotkowska ME, Tohge T, Fernie AR, Xue GP, Balazadeh S, Mueller-Roeber B**. **2015**. The Arabidopsis transcription factor MYB112 promotes anthocyanin formation during salinity and under high light stress. *Plant Physiology* **169**: 1862–1880.

**Mahmood K, El-Kereamy A, Kim SH, Nambara E, Rothstein SJ**. **2016a**. ANAC032 positively regulates age-dependent and stress-induced senescence in arabidopsis thaliana. *Plant and Cell Physiology* **57**: 2029–2046.

**Mahmood K, Xu Z, El-Kereamy A, Casaretto JA, Rothstein SJ**. **2016b**. The Arabidopsis transcription factor ANAC032 represses anthocyanin biosynthesis in response to high sucrose and oxidative and abiotic stresses. *Frontiers in Plant Science* **7**.

**Maki H, Sakaoka S, Itaya T, Suzuki T, Mabuchi K, Amabe T, Suzuki N, Higashiyama T, Tada Y, Nakagawa T, *et al.*** **2019**. ANAC032 regulates root growth through the MYB30 gene regulatory network. *Scientific Reports* **9**.

**Schweighofer A, Hirt H, Meskiene I**. **2004**. Plant PP2C phosphatases: Emerging functions in stress signaling. *Trends in Plant Science* **9**: 236–243.

**Shang J-Y, Lu Y-J, Cai X-W, Su Y-N, Feng C, Li L, Chen S, He X-J**. **2021**. COMPASS functions as a module of the INO80 chromatin remodeling complex to mediate histone H3K4 methylation in Arabidopsis. *The Plant cell* **33**: 3250–3271.

**Slawinski L, Israel A, Artault C, Thibault F, Atanassova R, Laloi M, Dédaldéchamp F**. **2021**. Responsiveness of Early Response to Dehydration Six-Like transporter genes to water deficit in Arabidopsis thaliana leaves. *Frontiers in Plant Science* **12**.

**Sun L, Zhang P, Wang R, Wan J, Ju Q, Rothstein SJ, Xu J**. **2019**. The SNAC-A transcription factor ANAC032 reprograms metabolism in Arabidopsis. *Plant and Cell Physiology* **60**: 999–1010.

**Takasaki H, Maruyama K, Takahashi F, Fujita M, Yoshida T, Nakashima K, Myouga F, Toyooka K, Yamaguchi-Shinozaki K, Shinozaki K**. **2015**. SNAC-As, stress-responsive NAC transcription factors, mediate ABA-inducible leaf senescence. *Plant Journal* **84**: 1114–1123.

**Thomashow MF**. **1999**. Plant cold acclimation: freezing tolerance genes and regulatory mechanisms. *Annu. Rev. Plant Physiol. Plant Mol. Biol* **50**: 571–99.

**Yang A, Dai X, Zhang WH**. **2012**. A R2R3-type MYB gene, OsMYB2, is involved in salt, cold, and dehydration tolerance in rice. *Journal of Experimental Botany* **63**: 2541–2556.
